# Keratose sponges in ancient carbonates – a problem of interpretation

**DOI:** 10.1101/2022.03.23.485445

**Authors:** Fritz Neuweiler, Stephen Kershaw, Frédéric Boulvain, Michał Matysik, Consuelo Sendino, Mark McMenamin, Rachel Wood

## Abstract

Increasing current interest in sponge fossils includes numerous reports of diverse vermicular and peloidal structures interpreted as keratose sponges in Neoproterozoic to Mesozoic carbonates and in various open marine to peritidal and restricted settings. Reports of their occurrence are fundamental and far-reaching for understanding microfacies and diagenesis where they occur; and fossil biotic assemblages, as well as wider aspects of origins of animals, sponge evolution/ecology and the systemic recovery from mass extinctions. Keratose sponges: 1) have elaborate spongin skeletons but no spicules, thus lack mineral parts and therefore have poor preservation potential so that determining their presence in rocks requires interpretation; and 2) are presented in publications as interpreted fossil structures almost entirely in two-dimensional (thin section) studies, where structures claimed as sponges comprise diverse layered, network, particulate and amalgamated fabrics involving calcite sparite in a micritic groundmass. There is no verification of sponges in these cases and almost all of them can be otherwise explained; some are certainly not correctly identified. The diversity of structures seen in thin sections may be reinterpreted to include: a) meiofaunal activity; b) layered, possibly microbial (spongiostromate) accretion; c) sedimentary peloidal to clotted micrites; d) fluid escape and capture resulting in birdseye to vuggy porosities; and e) molds of siliceous sponge spicules. Without confirmation of keratose sponges in ancient carbonates, interpretations of their role in ancient carbonate systems, including facies directly after mass extinctions, are unsafe, and alternative explanations for such structures should be considered. This study calls for greater critical appraisal of evidence, to seek confirmation or not, of keratose sponge presence. (259/300 max, for Sedimentology)

## INTRODUCTION AND AIM

This study addresses the issue of recognition of keratose sponges in thin sections of carbonate rocks, important because claims of their preservation potentially extends the body fossil record deep into the Neoproterozoic (Turner, 2021), thereby affecting analysis of sedimentary facies containing these structures across a long time range. For the purposes of our investigation, sponges present two fundamental forms that require understanding in relation to their preservation as fossils: 1) those with mineral spiculate skeletons (Figs 1E, F, 2A, B, 3, 4A-D) versus 2) those lacking such mineral parts (Fig. 1A-D, 2C, D, 4E-F), the latter constituting the Keratosa group of demosponges (Fig. 5). The taxonomic status of the best-known fossil aspiculate sponge *Vauxia*, examples of which are shown in Fig. 4E-F, is unconfirmed (Ehrlich *et al*., 2013). Sponges with mineral spiculate skeletons are most easily studied in hand specimens (e.g. Botting *et al*., 2017, Rigby *et al*., 2008) as either whole fossils, or as disaggregated spicules, noting that on death sponges generally break up very quickly, spicules readily dissolve and normally disappear to leave no record (Debrenne, 1999; Wulff, 2016). Thus, knowledge of the geological history of sponges has relied greatly on molecular clock phylogeny (e.g. Schuster *et al*. 2018; Kenny *et al*. 2020), see Fig 5. Fossil sponges with mineral skeletons, where preserved, are thus relatively easily recognizable in hand specimens but in thin sections are open to some interpretation, particularly if disaggregated (see Flügel, 2004, p. 495, 799). However, keratose sponges are significantly more problematic, yet have been inferred in a range of facies in the rock record, addressed next.

**Figure 1.**
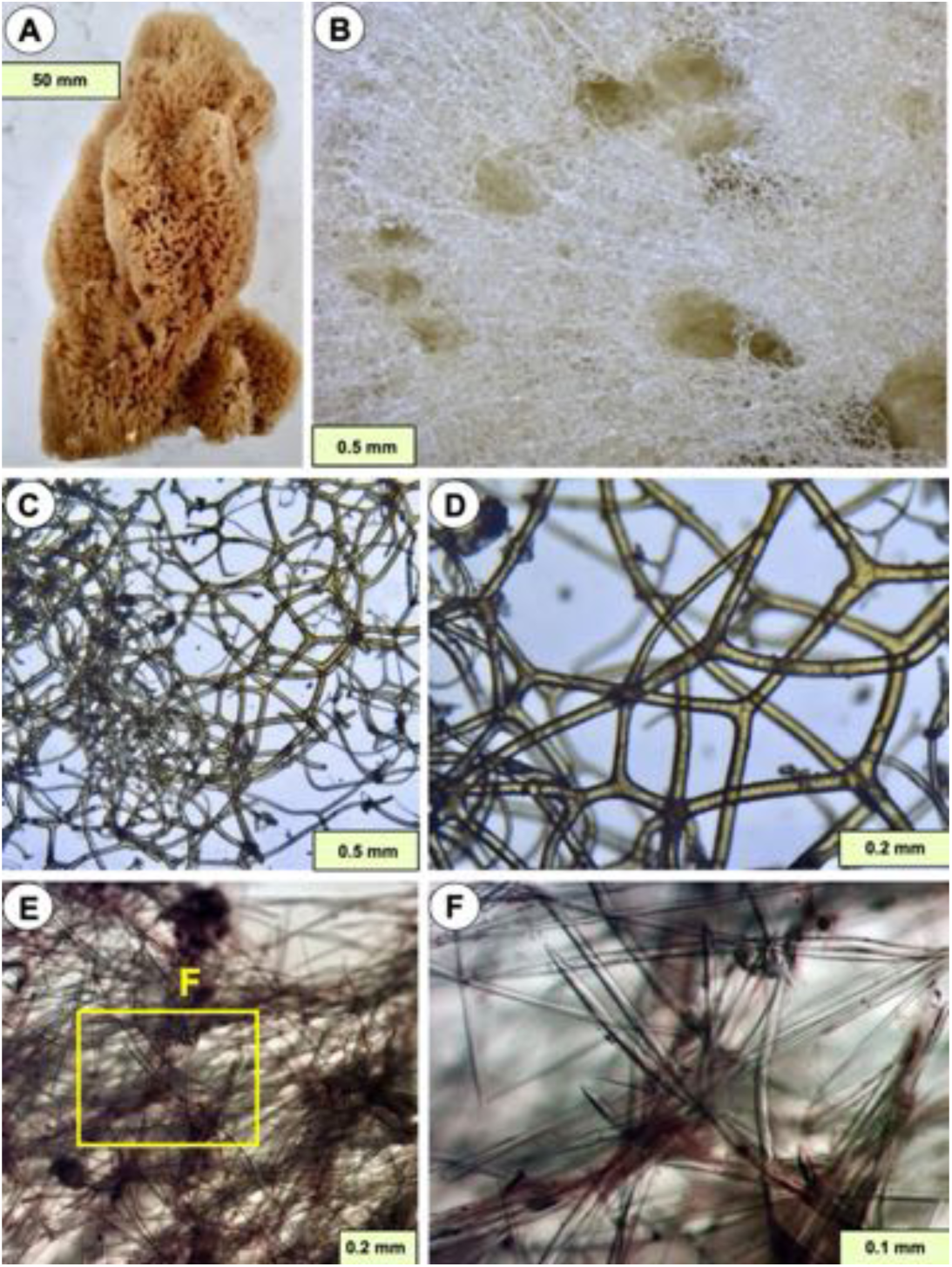
Examples of modern sponges showing contrasting construction of spiculate and keratose sponges. (A) Side view of a modern commercial keratose sponge consisting of only spongin (all the soft tissue is removed) showing its form is maintained by the spongin network. (B) Detail of sponge surface showing the spongin network and oscula (large holes) accommodating the excurrent canal system, a feature missing in the reported cases of fossil interpreted keratose sponges. (C-D) Details of the branched nature of the spongin network in A, showing branches and curved features; note that if ancient carbonate structures illustrated in this study, and references herein, represent keratose sponges, then the spongin networks shown in these two photographs would need to be preserved as calcite and the intervening empty space occupied by micrite. (E-F) Details of spiculate structure of *Phakellia robusta* Bowerbank, Shetland, Scotland. Bowerbank collection, Natural History Museum, London, sample, NHMUK 1877.5.21.420.

**Figure 2.**
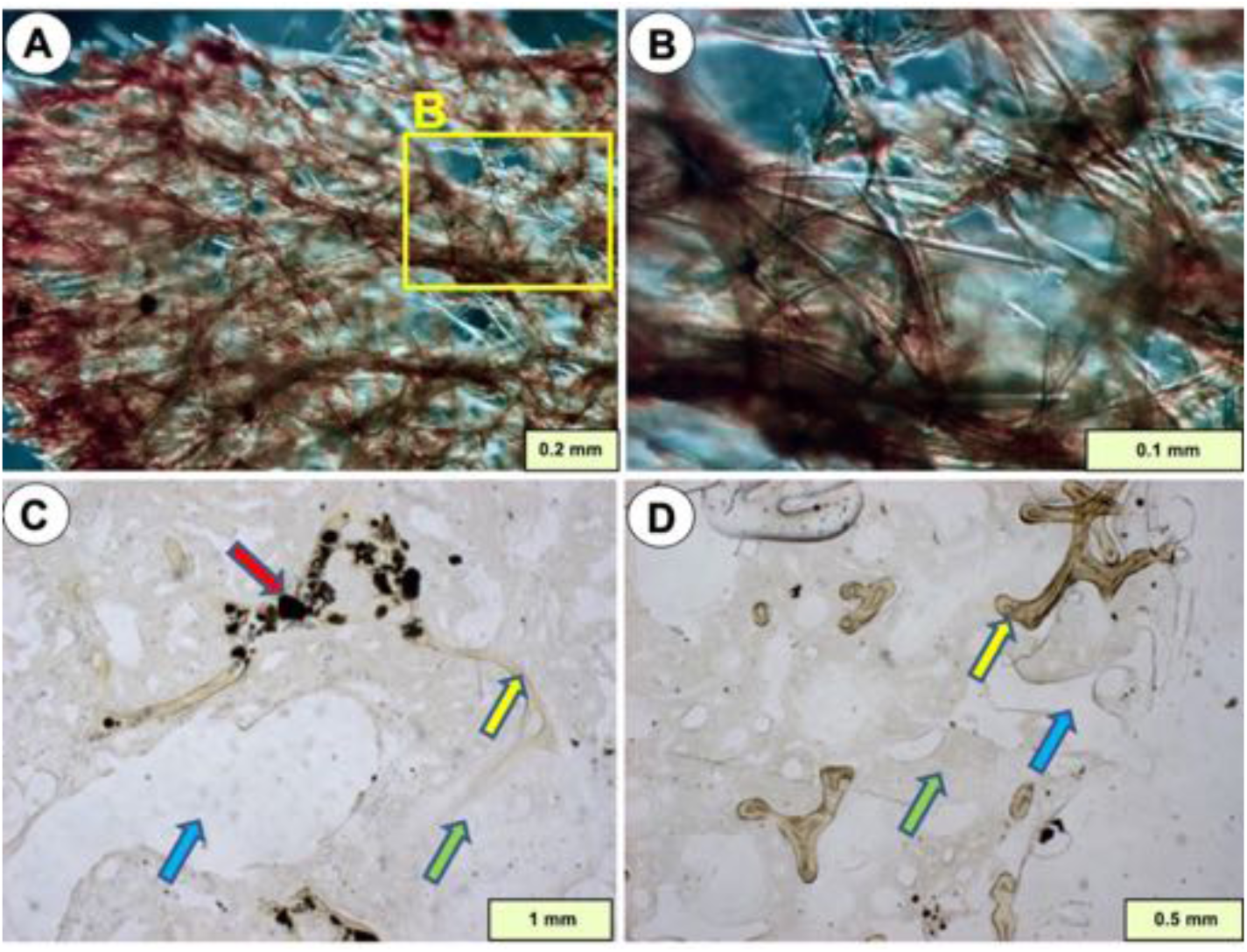
Examples of two modern sponges demonstrating more diversity of sponge architecture relevant to this study. (A, B) *Axinella verrucosa* [should be Esper but the label appears to say “Schmidt”], from the Adriatic Sea, showing spongin fibres encrusted by spicules; if the spongin component is to be preserved as a fossil, then it may be expected that sponges comprising both spongin and spicules should show both components in the fossil record, and this has not yet been demonstrated, see text for discussion. Bowerbank collection, Natural History Museum, London, sample NHMUK 1877.5.21.1239. (C, D) *Ircinia* keratose sponge showing strong primary spongin fibres (yellow arrow), soft porous mesohyl tissue (green arrow) with water canals (blue arrow), and incorporated detrital particles (red arrow, common in sponges). Joulters Cay, Bahamas.

**Figure 3.**
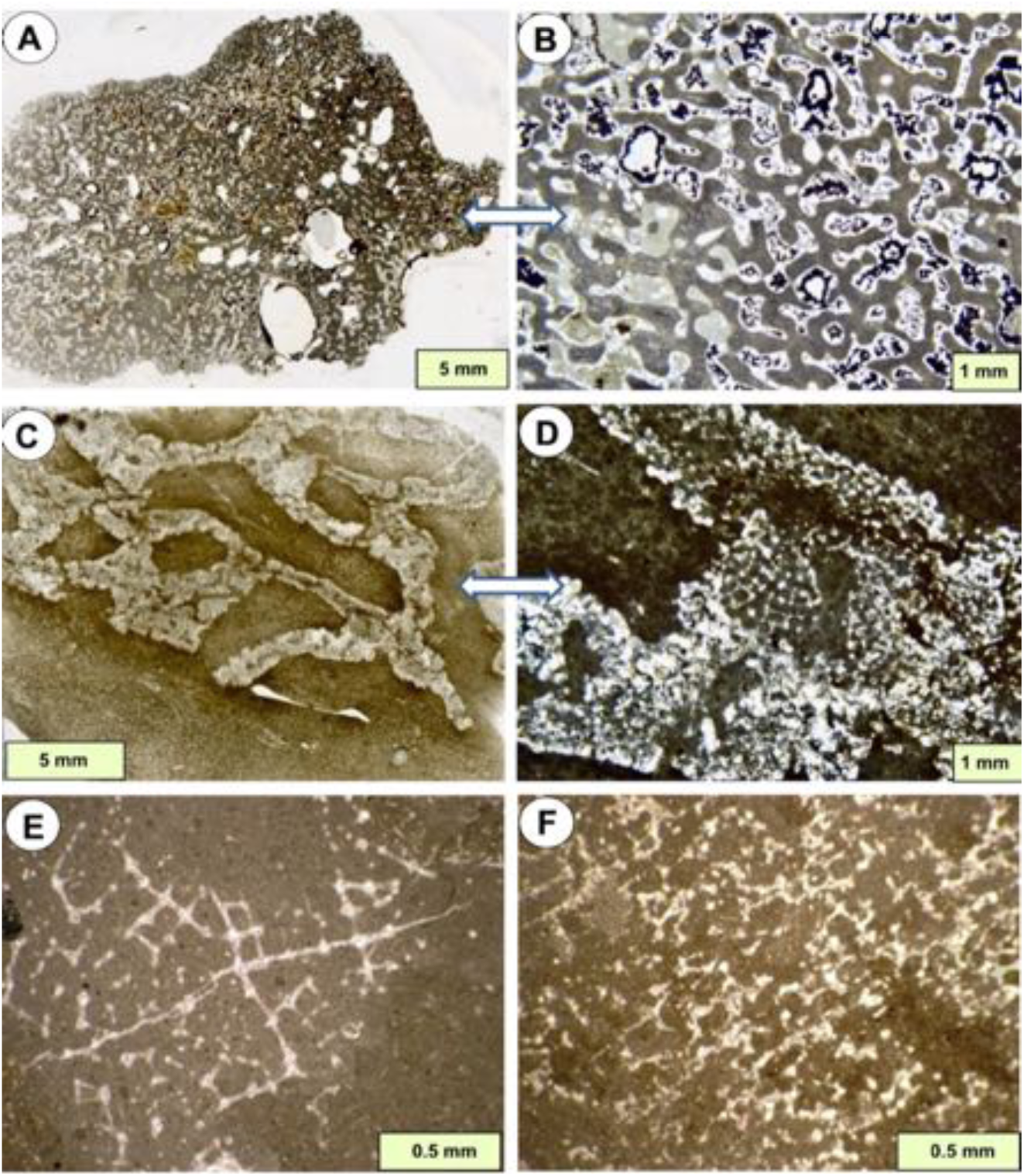
Examples of sponge mummies. (A, B) Calcified Cretaceous sponge from the Faringdon gravels, England, showing the sponge structure preserved as calcite and infilled with micrite. (C, D) Calcified Cretaceous sponge from the Chalk Group, Beachy Head, Eastbourne, England, showing a partially preserved spiculate network. (E) Hexactinellid sponge mummy showing details of spicule network preservation as calcite. (F) Lithistid sponge showing desma spicules preserved in calcite. E and F from Dalichai Formation, Bajocian-Callovian (Jurassic), Alborz Mountains, northern Iran. Photographs kindly provided by Andrej Pisera.

**Figure 4.**
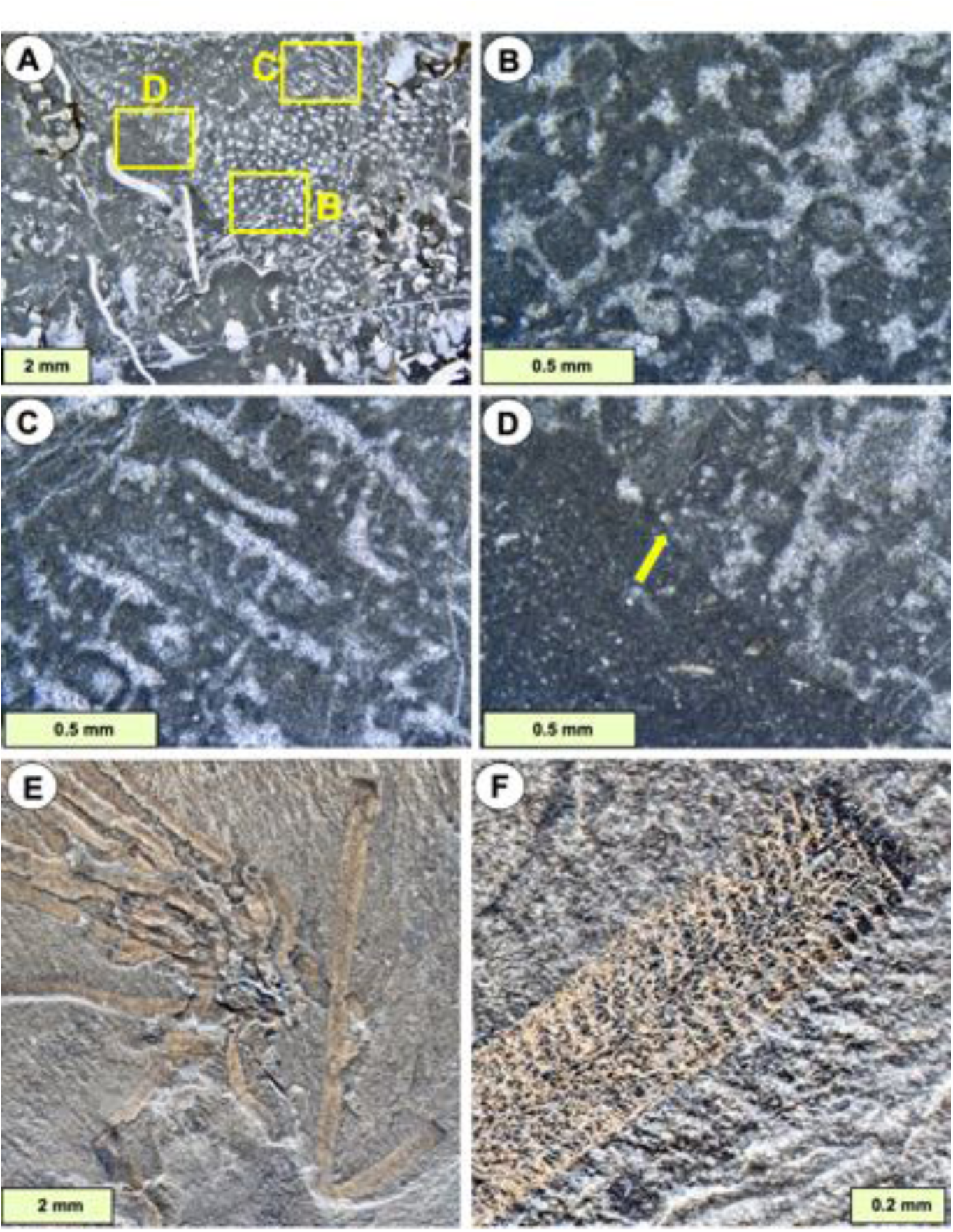
(A-D) Calcified spiculate sponges, may be lithistids, showing the rectilinear network structure described in the text. (A) the sponge forms a discrete object in the upper right half, locations of B-D are indicated. (B-C) Details of transverse (B) and vertical (C) sections of spiculate structure, noting spicules are preserved as calcite. (D) Detail of margin of sponge showing its sharp contact (arrow) with surrounding micrite. Church Reef, Filimore Formation, L. Ordovician, Utah. (E, F) The aspiculate fossil sponge *Vauxia gracilenta* Walcott, 1920 from the Burgess Shale, regarded as one of the best examples of aspiculate, possibly Keratose sponges, see Walcott (1917). Specimen NHMUK PI S3071 in the Natural History Museum, London.

**Figure 5.**
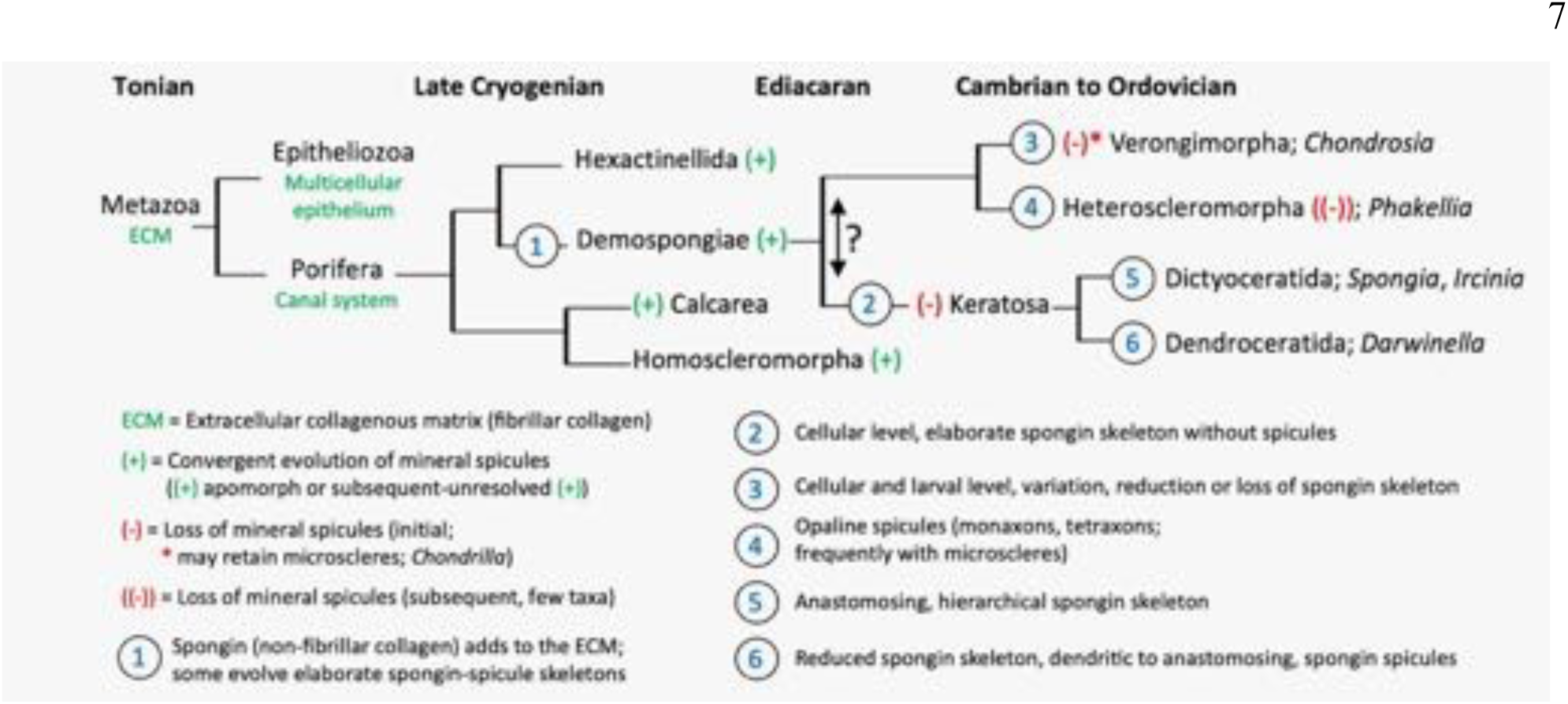
Summary evolutionary history of sponges (see Schuster *et al*. 2018; Kenny *et al*. 2020), drawing attention to major events including the origin of the Keratosa, that are the principal subject of this paper.

The origin of interpretations of fossil keratose sponges in carbonates seems to have been a study by Szulc (1997) who described stromatolites from restricted lagoonal facies (Matysik, 2016) in the Middle Triassic Muschelkalk carbonates from Upper Silesia; Szulc inferred that pockets and layers of porous micrite within the stromatolites are sponges (Fig. 6).

**Figure 6.**
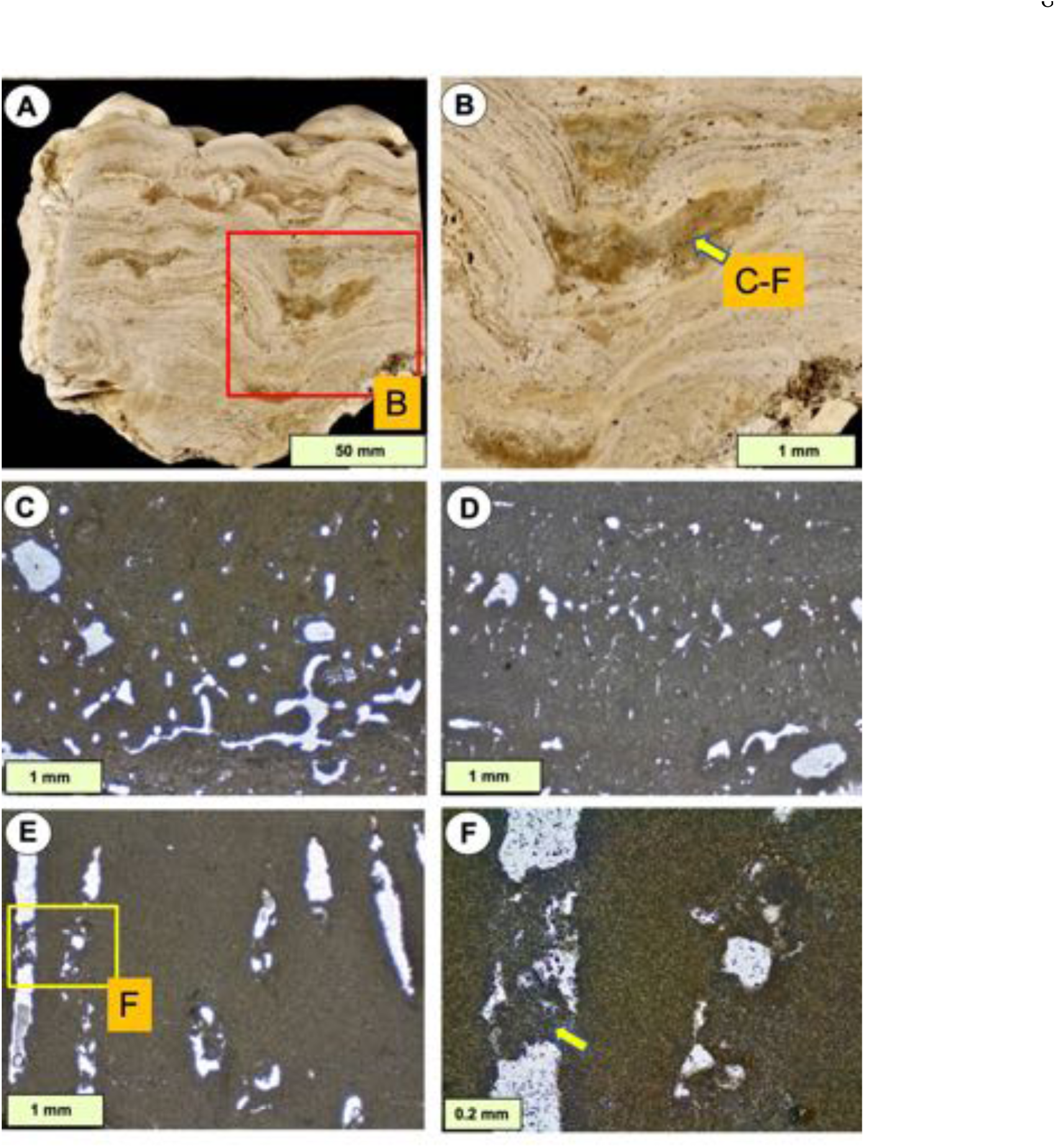
Stromatolite samples proposed by Szulc (1997) to contain sponges. (A, B) Vertical sections through stromatolite showing interlayered sediment in which the porous fabrics (yellow arrow) occur that were considered as sponge by Szulc. (C, D) Vertical thin section views of the interlayered sediment in A, B, showing organised porosity as subvertical voids. (E, F) Detail of one pore, showing partial infill with micrite (yellow arrow), evidence that the pore must have been open to the sea floor to collect sediment and thus not consistent with interpretation as a permineralised sponge. From the collections of Joachim Szulc; boundary between Diplora Beds and overlying Tarnowice Beds, Muschelkalk, Middle Triassic, Libiąż Quarry, Upper Silesia, Poland.

However, Szulc (1997, p. 14) did not give criteria for recognition of sponges, the porous vermicular structures he described have no spicules (Fig. 6) and are preserved in dolomite. Also there is an issue regarding the likelihood of occurrence of sponges in combination with stromatolites in restricted facies, not known in modern environments. Confusingly Szulc (1997) noted similar deposits from Thuringia that are silicified, from which he inferred the proposed sponges were siliceous, but without any supporting evidence. Non-spiculate sponges were similarly inferred by Reitner *et al*. (2001) and Reitner & Wörheide (2002, fig. 10) for Devonian mud-mounds. Then, in a landmark study, Luo & Reitner (2014) used serial grinding and imaging methods to construct 3D views, leading to their interpretation of possible fossil keratose sponges (aspiculate) to explain vermicular structure in other carbonates. However, their reconstructions were inconclusive, with their original interpretation unsubstantiated, as recognized by Luo & Reitner (2014) themselves, who used terms such as “most likely”, “putative” and “preliminary”.

Subsequently, Luo & Reitner’s (2014) work was developed by Luo & Reitner (2016), then these studies were used to support numerous claims of keratose sponges in other carbonates from Neoproterozoic to at least Triassic time (see compilation of publications in Table S1), without verification, and no others have attempted 3D reconstruction. Instead, evidence is presented in 2D images (thin sections), commonly at low-resolution where details are not clear, and rely on broad textural features as the basis of a sponge interpretation.

Thus the problem that keratose sponges present is lack of reliable identification of their fossilized remains in any part of the rock record; and there is no known diagenetic process that could transform the entity of the porous soft tissue (between the spongin fibres, see Fig. 2C, D) into the largely homogenous micrite that dominates interpreted keratosans. In contrast, Figs. 3 and 4 show the types of preservation of fossil sponges commonly encountered in thin section, that, with the exception of Fig. 4E & F, comprise mineral parts. Therefore, this study draws attention to the difficulties of understanding the fabrics of interpreted keratose sponges in carbonate rocks that may instead be viewed diversely as *Problematica* (definite fossils, the affinities of which are not known), fragments of altered siliceous sponges, graphoglyptid trace fossils or even dubiofossils. Thus, the aim of this study is to bring into focus the problem of recognition of keratose sponges, and consider alternatives that may be explored in future research. In order to explain the issues fully, a background review is provided of the issues around fossil sponges that necessarily involves description of relevant features of modern sponges. Then classification, description and discussion of the range of fabrics of carbonate rocks published as interpreted keratose sponges is presented, relevant to microfacies analysis of carbonate rocks. The focus is on four key settings in which keratose sponges have been interpreted: Neoproterozoic carbonates, consortia between stromatolites and sponges (especially Triassic), Cambro-Ordovician carbonates, and carbonate facies in the aftermath of mass extinctions.

## BACKGROUND

Ambiguities in the interpretation and thus classification of sedimentary carbonate materials are widespread; such ambiguities are mostly related to the structure and volumetric importance of relatively small microcrystalline grains (Lokier & Al Juanabi, 2016). Sources of error include the problems of identification, in thin-section, of structures that may be fossils, in terms of form, functional design and skeletal microstructure (Knoll, 2003; Flügel and Munnecke, 2010), thus constituting a grey zone between clearly identifiable and suspect structures. This grey zone comprises objects that may be considered in three types: a) biogenic but require interpretation in terms of basic taxonomic placement (*Problematica sensu lato*; Jenner & Littlewood, 2008; e.g. Paleozoic *Halysis* as red alga, cyanobacteria, green alga or tabulate coral; Zheng *et al*., 2020), b) distinct structures but inconclusive in terms of biogenicity (dubiofossils of Hofmann, 1972), and c) distinct structures that are certainly abiotic in nature (pseudofossils, full discussion in McMahon *et al*., 2021). Another aspect is that the granularity of carbonate deposits does not necessarily relate to sedimentary processes; it might be post-depositional in nature due to meio-to endofaunal activity, localized microburrow nests or even diagenesis (Debrenne *et al*., 1989; Wood *et al*., 1993; Pemberton & Gingras, 2005; Löhr & Kennedy, 2015; McMenamin, 2016; Wright & Barnett, 2020). Furthermore, this grey zone applies to cases that extend into deep time and even touches exobiology (Cloud, 1973; e.g., biogenicity criteria for tubular filaments and lamination; chemical gardens comprising inorganic processes resulting in structures resembling organisms; molar-tooth structures; see Grotzinger & Rothman, 1996; Awramik & Grey, 2005; McMahon *et al*., 2017, 2021; McMahon & Cosmidis, 2021). This range of fabrics persists throughout the Phanerozoic in various ways; examples are: the biogenicity of stromatactis and lamination (Bathurst, 1982; Bourque & Boulvain,1993; Awramik & Grey, 2005; McMahon *et al*., 2021); the formation of peloids (Macintyre, 1985); the significance of the polymud fabric (Lees & Miller, 1995; Neuweiler *et al*., 2009); or some drag marks, *Rutgersella* and *Frutexites* (Cloud, 1973; Retallack, 2015; McMahon *et al*., 2021).

During the last decade, molecular phylogenetic studies have shed new light on the traditional taxonomic and phylogenetic framework of sponges, revealing or confirming several polyphyletic groups, establishing new clades, and constraining respective divergence-time estimates (Gazave *et al*., 2012; Erpenbeck *et al*., 2012; Morrow & Cárdenas, 2015; Schuster *et al*., 2018; Kenny *et al*., 2020), see Fig. 5. A valuable general reference for sponge groups is de Voogd *et al*. (2022). Spongin is considered to have evolved in tight connection within the demosponge lineage (Morrow & Cárdenas, 2015). Sponge spicules may not represent an essential character of early sponge evolution (Ax, 1996), and were secondarily lost in a multiple and convergent manner (Fig. 5). Keratose sponges are distinguished from other aspiculate demosponges (Verongimorpha, some Heteroscleromorpha according to Erpenbeck *et al*., 2012) at the cellular level in combination with the details or even absence of an elaborate spongin skeleton (Erpenbeck *et al*., 2012).

As indicated in the Introduction, the secure identification of fossil sponges essentially relies on spicules, commonly identified according to their specific design and arrangement, comprising: form, orientation; assemblage and mineralogy (examples in Figs 1-4). Sponge form may also be preserved *via* a process referred to as mummification, that is early calcification (thus lithification) of the sponge tissue with its associated sediment, to preserve the sponge shape and organisation sufficiently enough to allow recognition as a sponge (canal system, preservation of non-rigid spicular architecture; Fritz, 1958; Bourque & Gignac, 1983; Reitner & Keupp, 1991; Pisera, 1997; Neuweiler *et al*., 1999; Reitner & Wörheide, 2002; Neuweiler *et al*., 2007 with references therein). In other cases, there are specific secondary calcareous skeletons that leave a good record, preserved as, e.g., stromatoporoids, chaetetids, inozoans and sphinctozoans, at least one of which (*Vaceletia*) is considered a coralline keratose sponge (Wörheide, 2008). Some sponges leave distinct ichnofossils (*Entobia*), that may contain spicule evidence of their formation (Reitner & Keupp, 1991, Bromley & Schönberg, 2008). Biomarkers might be of additional value (e.g. Love *et al*. 2009; see also Antcliffe *et al*., 2014), but their study requires a detailed understanding of both history of fluid flow and molecular analogues of possible other origin. Confusingly, some foraminifera use sponge spicules to agglutinate their tests (Ruetzler & Richardson, 1996; Kamenskaya *et al*., 2015) and thus need careful study to distinguish them from sponges.

Against the background of well-known modern aspiculate sponges (Keratosa and Verongimorpha) with their enormous architectural variability (Manconi *et al*., 2013; Stocchino *et al*., 2021), the situation is naturally precarious for claims of fossil non-coralline keratose demosponges to be preserved in limestones and dolostones. The body shape stability of keratose sponges (Fig. 1A, B) relies on fibrillar collagen as a key component of their extracellular collagenous matrix (ECM), that at micro-to macroscale is combined with a highly elastic and elaborate organic skeleton composed of the non-fibrillar collagen spongin (Exposito *et al*., 1991; Erpenbeck *et al*., 2012; Ehrlich, 2019). Indeed, the fossil record of non-spiculate sponges was described by Reitner & Wörheide (2002) as being poor, noting that the vauxiid sponges of the middle Cambrian Burgess Shale are the best examples (Fig. 4E, F). However, Ehrlich *et al*. (2013) revealed that those sponges contain chitin (as other sponges and a number of invertebrates do), but not spongin. Fan *et al*. (2021) classified aspiculate vauxiid sponges in the Chengjiang biota (early Cambrian) as keratose sponges, preserved in partly silicified form.

Apart from vauxiids noted above, in literature search no verified cases of keratose sponges have been found in the entire rock record. An important aspect is that, within modern spiculate sponges, there are variable amounts of non-spicular material in proportion to the spicule content. Thus, it is necessary to appreciate the detailed features of sponges with and without spicules. Fig. 2A, B shows details of a modern heteroscleromorph spiculate demosponge that has spongin fibres encrusted with opaline spicules, and is one example of the common occurrence of tightly connected spicules and spongin fibres known for many decades (Axinellidae Carter, 1875), yet there is no report of a respective thin-section fossil that replicates such a distinct composite skeletal architecture. Singular claims for fossil Axinellidae are unconfirmed (Reitner & Wörheide, 2002, their Fig. 9, which may instead be spicule-preserving *Entobia*). In addition there are modern non-spiculate (keratose) sponges comprising conspicuous primary fibres of spongin up to 250 µm thick and distinctively cored by sand grains (Irciniidae, Gray 1867; see also Manconi *et al*., 2013) (Fig. 2C, D) but again, there is no fossil record. For further comparison, Figs 3 and 4A-D demonstrate carbonate fabrics typical of preserved spicule-bearing sponges, but in these there is no indication of the spongin component that is presumed lost in decay and diagenesis. Such details are important to gain an understanding of how such soft-tissue structures might be preserved, and comparisons between these and fossil cases are made later in this paper. Nevertheless, prominent examples of interpreted keratose sponges are in studies by Luo & Reitner (2016), Lee & Hong (2019), Lee & Riding (2021a, b), Baud *et al*. (2021), Pei *et al*. (2021), Pham *et al*. (2021), Gischler *et al*. (2021) and Turner (2021). None of those studies highlighted biostratinomy in combination with porosity evolution and diagenesis. Criteria for distinguishing, for example, carbonate microfabrics attributed to sponges from microbial deposits in ancient carbonates (Wallace *et al.,* 2014; Shen & Neuweiler, 2018) are not defined, and there are numerous other possible interpretations that are explored in this study.

## MATERIAL AND METHODS

In order to address the ideas regarding keratose sponges in the carbonate rock record, a range of samples from was used: Cambrian of North China, Cambro-Ordovician of Nevada and Utah, Silurian of south China, Devonian of Belgium, Viséan of Boulonnais region (France) and Triassic of the Upper Silesian region (Poland) including some original samples from the Triassic material from Poland and Israel used by Szulc (1997) and Luo & Reitner (2014, 2016). For basic reference, examination was made of the original spongiostromate material (Visé Group, Namur region) of Gürich (1906) stored at the Royal Belgian Institute of Natural Sciences (Brussels) and a selection of modern spiculate and non-spiculate sponges illustrated by: Bowerbank (1862) stored at the Natural History Museum London (UK); and from personal collections of the authors. Some published figures are reproduced under Creative Commons licences. Polished rock samples and thin sections were studied under plane-polarised (PPL) and cross-polarised (XPL) light, supplemented with selected cathodoluminescence (CL) and UV fluorescence views.

## RESULTS

In an attempt to follow the history of this topic, examination was made of Szulc’s (1997) sample material of stromatolitic deposits stored in the Jagellonian University, Poland, illustrated here with new thin sections and cathodoluminescence; then the study was developed to address other structures claimed as keratose sponges. Perusal of the literature and primary material led to recognition that structures interpreted as fossil keratose sponge may be divided into five broad fabric types differing in context, architecture and microstructure. Some overlap occurs between the five categories, thus some examples presented here may fit into more than one type. All are based on two-dimensional views in thin sections. All are composed of areas of sparite intermingled with micritic material, the latter commonly comprising homogenous micrite, but in some cases showing clotted to granular fabrics. In some other cases reported in literature, they occur within shells but not in the matrix surrounding the shells (Park *et al*., 2017, fig. 3); in still other cases they occur in discrete patches in micrite, and may have been burrows (Park *et al*., 2015, fig. 4D); these two cases may reflect re-burrowing of organic-rich sediment encased in pre-existing burrows and shells. Many other examples occur within early-formed cavities in early-lithified limestone (Lee *et al*., 2014). Some of the micritic material contains fossils (e.g. Lee & Hong, 2019, fig. 2c) which are difficult to explain if these were keratose sponges.

In Figs 6-17, according to the ideas of keratose sponge interpretation, the curved and irregular sparite patches represent the position of the original spongin structure and the micrite infill represents where the sponge soft tissue was located. Clearly, of great importance is to explain how: a) keratosan sponges could be preserved through a biostratinomic and diagenetic process that began with an elaborate organic skeleton made of spongin enveloped by a canal-bearing soft tissue and ended with sparitic calcite in a microcrystalline groundmass that comprises these fabrics; and b) if spongin components are present in fossils, why are they not visible along with spicule remains in spiculate sponges in at least some cases (c.f. Fig. 2A,B)?

### Layered fabrics

Within the Triassic stromatolites regarded by Szulc (1997) as containing sponges (Figs 6, 7), the possible sponge component forms faint to prominent micrite layers containing porous network fabrics. Luo & Reitner (2014) used material from the same horizons.

**Figure 7.**
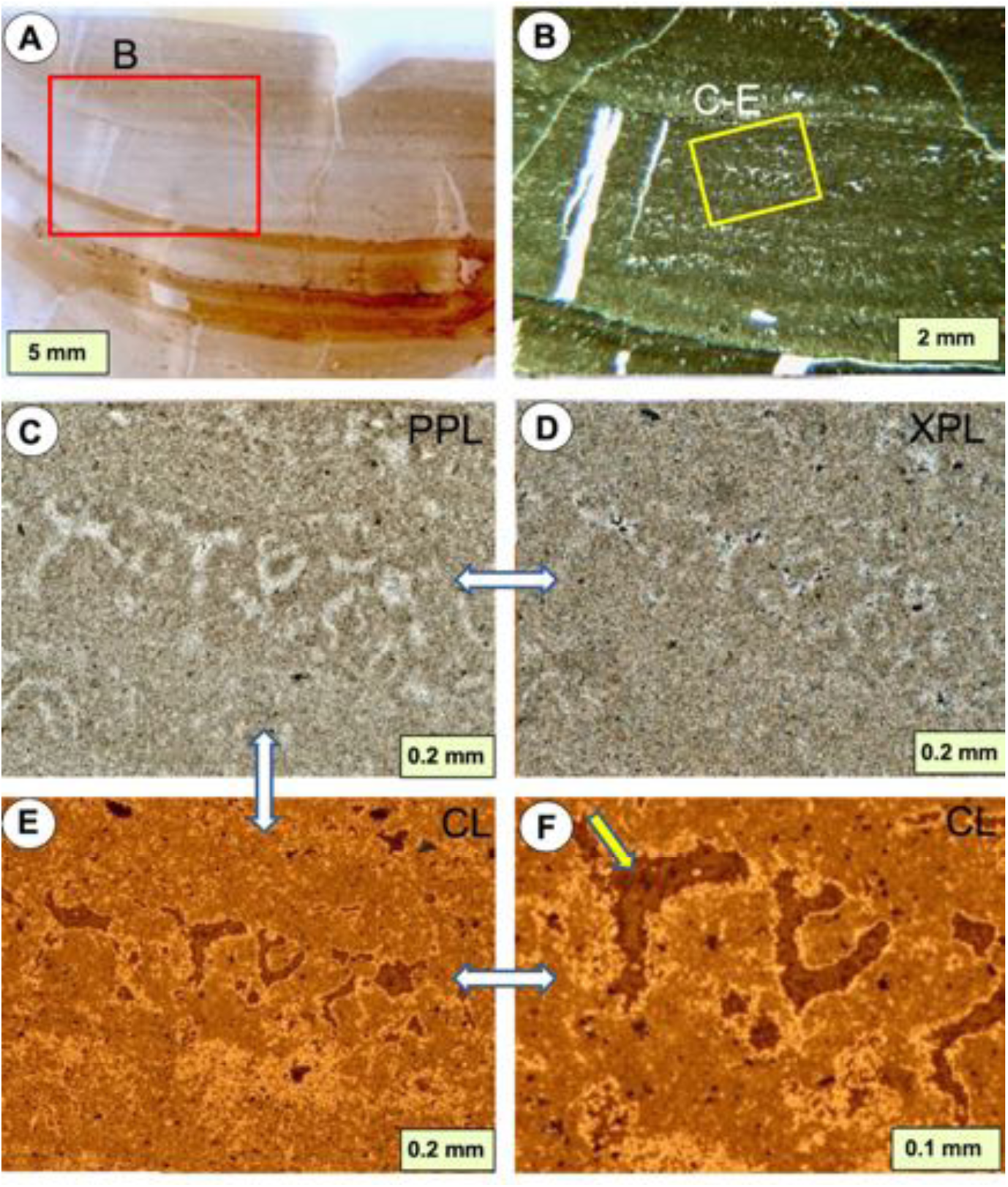
Vertical sections from a stromatolitic horizon, containing vermicular structures. (A) Whole thin section, showing prominent layered structure. (B) Detail of box in A, showing locations of C-F. (C, D) PPL (C) and XPL (D) views of vermicular structure within the stromatolite layers, showing sparite cement in the light areas. (E, F) CL view of C and D, with enlargement in F, showing a sequence of dull to no luminescence in the sparite areas (arrow), while the micrite areas contain a mixture of bright and dull luminescence. Note that the edges of the micrite against the sparite shows a higher degree of brighter luminescence. This pattern is interpreted to indicate that the sparite areas were vacated and infilled with cements, thus showing difference in diagenetic history from the micrite. From the collections of Joachim Szulc, sampled by him from the Ladinian (late Middle Triassic), Negev area, Israel.

Layers in these materials broadly match the concept of spongiostromates, introduced by Gürich (1906) to convey their open architecture and layered bioaccretionary character (Figs 7, 8). The spongiostromate microstructure represents a microporous fabric of likely microbiotic accretion, generally blurred, grumelous to peloidal, at best faintly tubular to cellular/vesicular. This is in opposition to the porostromate microstructure that displays well-defined micro-organismic outlines preserved in growth position (Monty, 1981; for comparison, see Turner *et al*., 2000; Flügel, 2004, p. 122). In this context layers comprise micrite normally enclosing somewhat irregular areas of sparite (Fig. 7), although some have open pores and others with micrite in the pores (e.g. Fig. 6F); the micrite may also include small bioclasts. Spongiostromate structures present a problem of interpretation because they have a spongy-looking fabric, in the common-English understanding of the term sponge, but without indication of a biological sponge nature, noting that Gürich’s (1906) monograph illustrations obviously do not show sponges. The oldest known occurrences of spongiostromate structures forming part of oncoids reach back to the Palaeoproterozoic (Schaefer *et al*., 2001; Gutzmer *et al*., 2002). Examples of spongiostromate-style fabrics interpreted to be keratose sponges may be seen in Luo & Reitner (2014, 2016), Pei *et al*. (2021a, b) and Lee & Riding (2021a, b). Stock & Sandberg (2019) illustrated layered fabrics of spongiostromate form in Devonian-Carboniferous boundary facies in Utah, which were presented as sponges, but their photographs are not sufficiently detailed to show structure that can be verified as sponge or not.

**Figure 8.**
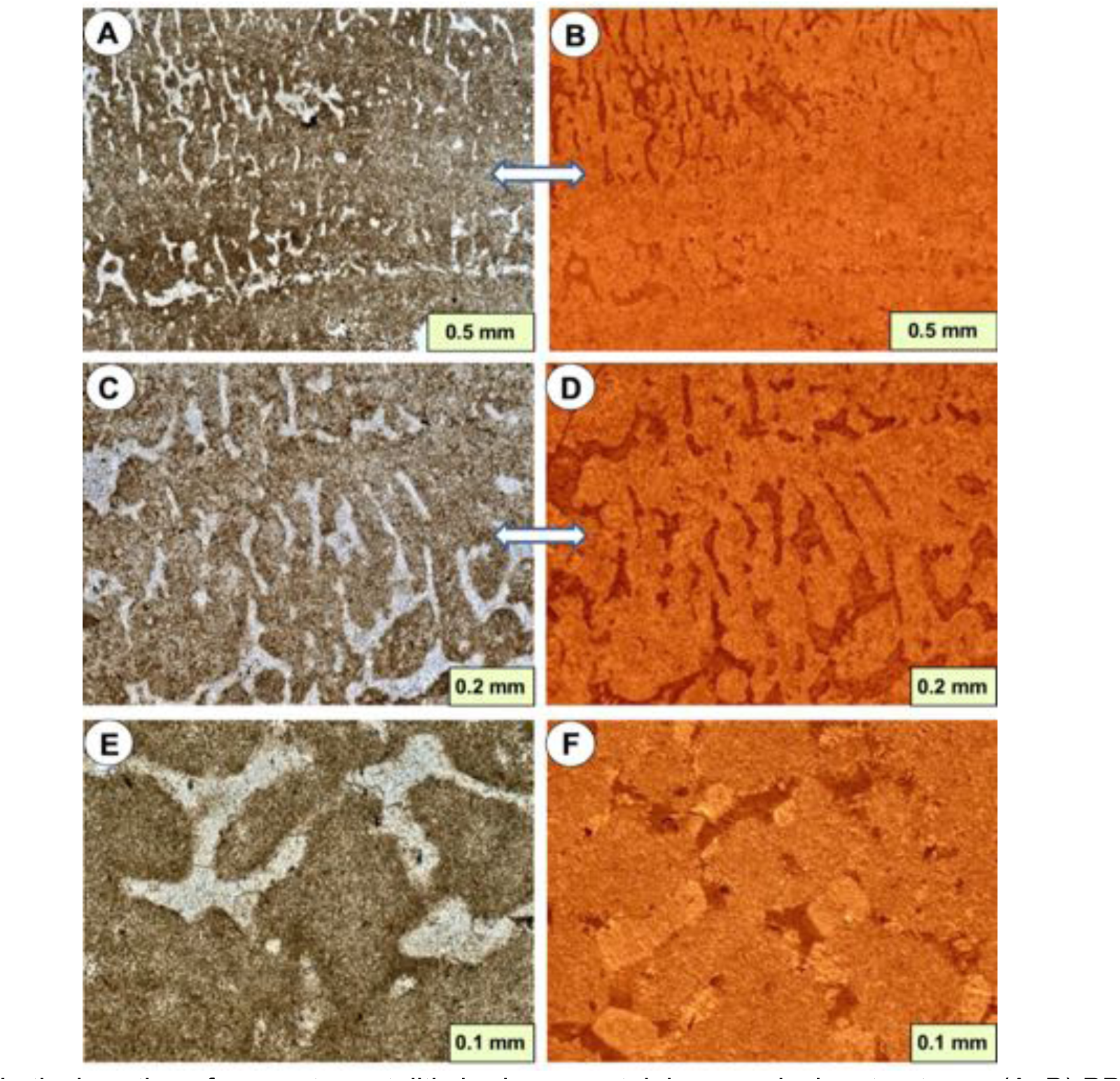
Vertical sections from a stromatolitic horizon, containing vermicular structures. (A, B) PPL (A) and CL (B) views of prominent vermicular structure in stromatolite. (C, D) Detail of structure from an adjacent area of thin section to A and B. (E, F) Detail of another area of this sample at greater enlargement. Images in this figure demonstrate the difference in CL response between the sparite and micrite areas, indicating their diagenetic histories are not coincidental. From the collections of Joachim Szulc, sampled by him from the Ladinian (late Middle Triassic), Negev area, Israel.

**Figure 9.**
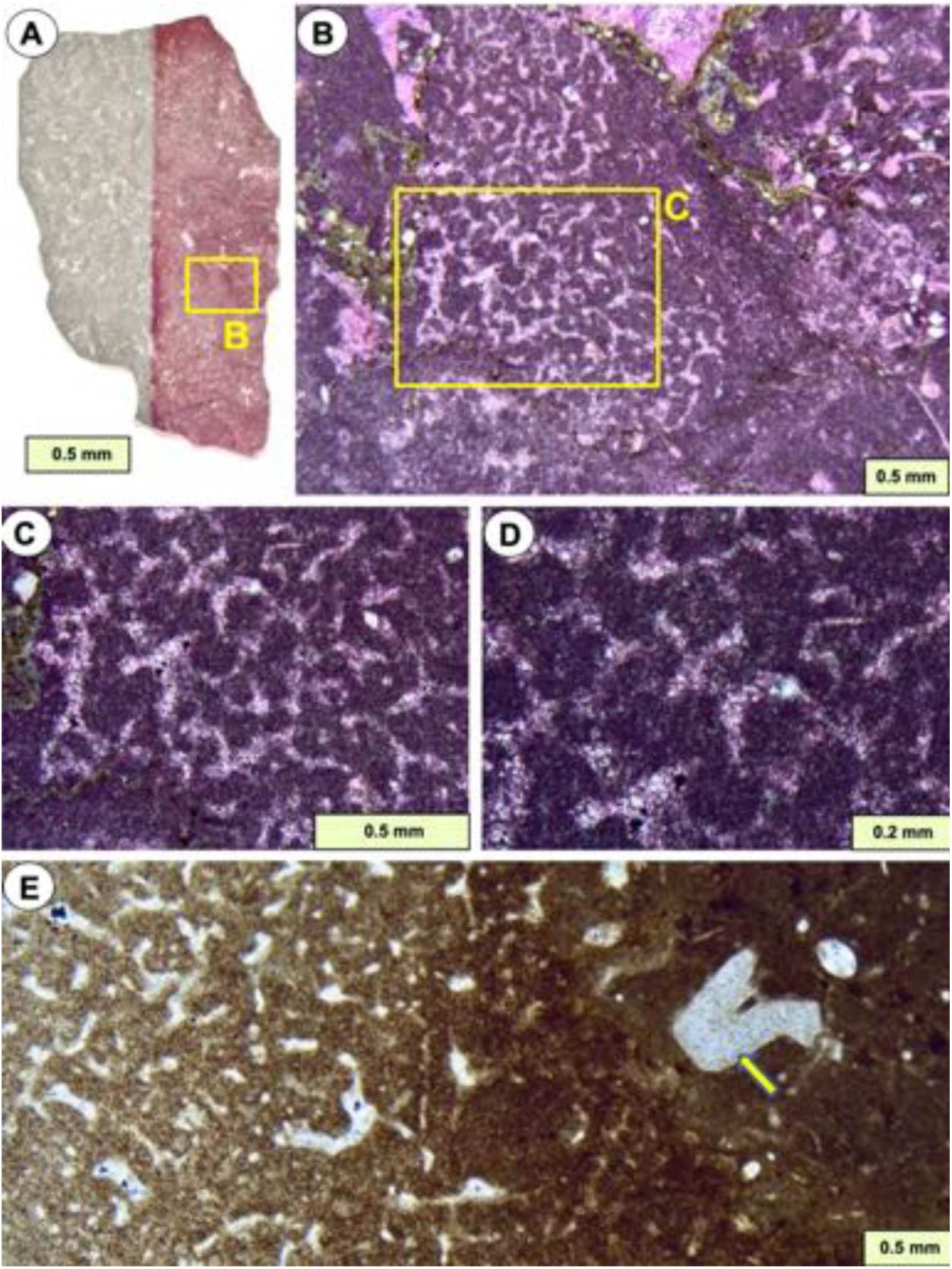
(A-D) Vertical sections through a small piece of limestone collected from matrix between stromatolite columns, showing a possible keratose sponge, although this is instead possibly a lithistid. (A) Whole thin section partly stained with ARS-KFeCN, showing a mottled fabric and the location of B. (B) The possible sponge forms a defined patch and shows a curved network of sparite-filled voids embedded in micrite. (C, D) Details of B (D is a detail of centre of C) showing the sparitic nature of the network preserved as red-stained (non-ferroan) calcite. A-D from Chalk Knolls, Notch Peak Formation, upper Cambrian, Utah. (E) Curved network of sparite in micrite, but with a diffuse margin; a crinoid columnal (arrow) is prominent in right hand part, outside the network. The network area may be a sponge, but its diffuse margin presents a problem of interpretation (Kershaw et al. 2021a). Huashitou reef, Ningqiang Formation, Telychian (lower Silurian), Guangyuan, northern Sichuan, China; specimen donated by Yue Li.

**Figure 10.**
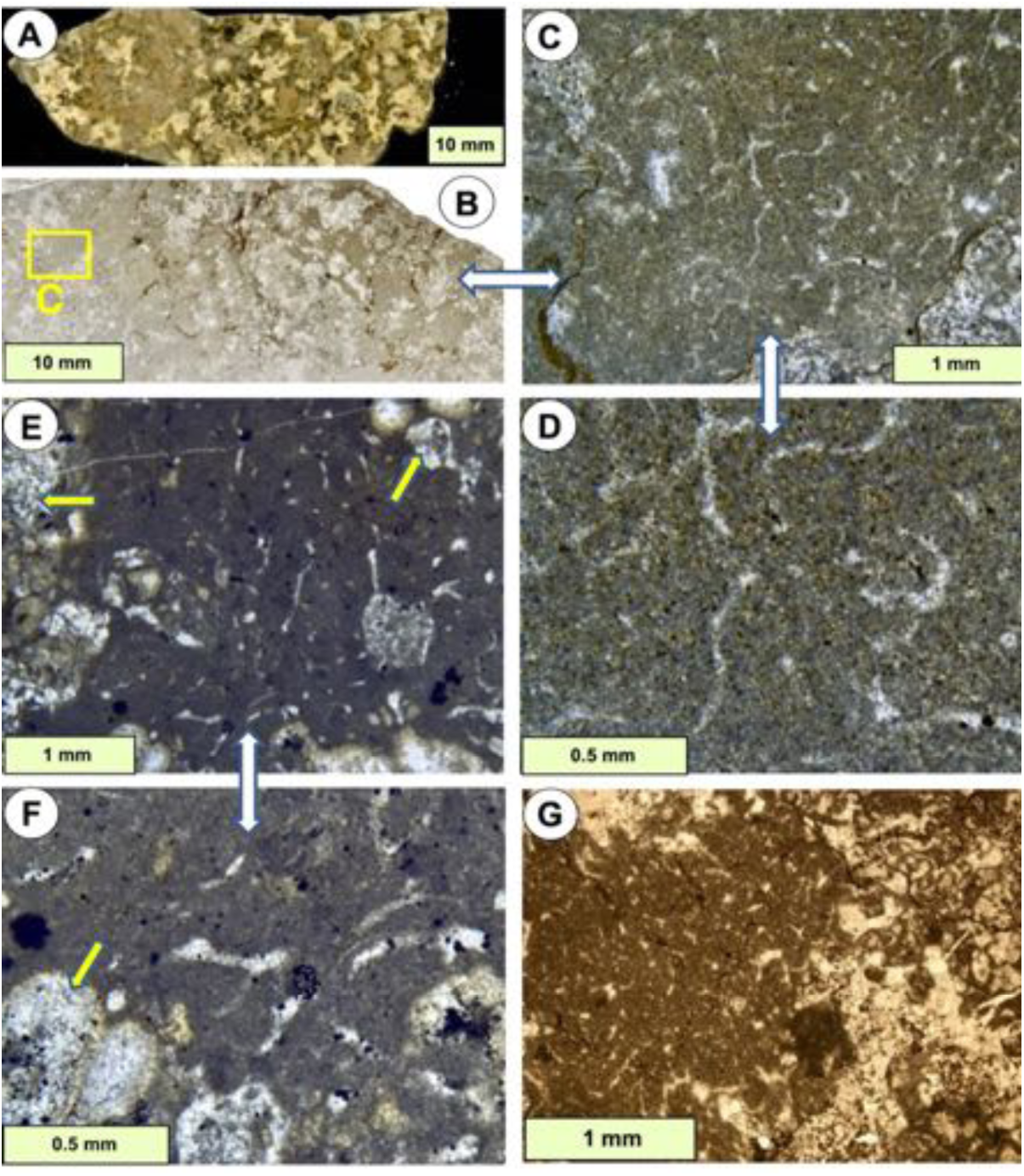
Examples of network fabrics in Permian-Triassic boundary microbialites from south China. (A, B) Hand specimen (A) and thin section (B) of microbialite a few cm above the end-Permian mass extinction horizon, showing recrystallised microbial calcite (lobate pale areas in B) with intervening micrite. (C, D) Enlargements of yellow box in B showing curved sparite areas in the micrite, that are of similar material to that interpreted as keratose sponge by Baud *et al*. (2021) and Wu *et al*. (2021). Permian-Triassic boundary interval, Baizhuyuan site, Huaying Mountains, Sichuan, China. (E, F) Another sample of microbialite after the end-Permian extinction, with curved sparite patches as in A-D, but in this case may comprise bioclasts. These two images also show lobate areas of light-coloured sparite in the edges (arrows), that are the calcimicrobe frame which constructured the microbialite (Kershaw *et al*., 2021b). Laolongdong site, Beibei, Chongqing, China. (G) The right-hand third of this view shows partially altered calcimicrobial structure, comparable to that described by Ezaki *et al*. (2008, Fig 8C from the Dongwan locality a few km along strike) as “spongelike”, mistakenly interpreted as sponges by some authors, see text for discussion. The left-hand two thirds show micrite infill, containing network fabrics, deposited between microbial branches. See text for discussion. Baizhuyuan site.

Layered fabrics may also show characters commonly described as birds-eyes, (connected) vugs and fenestrae, that are normally recognized as part of an intertidal to backreef carbonate system where degassing occurs in sediments exposed at the surface or in very shallow water (e.g. Tucker & Wright, 1990). They represent fabric-selective primary porosities with original voids commonly larger than the mean grain diameter. Laminae and sheets containing fabrics that may be reasonably interpreted as such, altogether being part of Triassic (Anisian) microbialites/stromatolites, were considered to be keratose sponges by Luo & Reitner (2014, 2016). In a subsequent step, the interpretation was developed to propose a distinction between a stromatolite and a sponge-microbial ‘consortium’ called keratolite by Lee & Riding (2021a). Conventionally, such structures are understood to form *via* the entrapment of gas bubbles, anhydrite precipitation and desiccation frequently in combination with dissolution and subsequent compaction in peritidal to intertidal (microbial) environments (Shinn, 1968). More recently, Bourillot *et al*. (2020) provided more details and a number of distinguishing parameters indicating how these porous microbialites/stromatolites may form their laminated-micritic, laminated-peloidal microfabrics.

Layered fabrics shown in Figs 7 and 8 compare plane light views with CL in paired images; the CL views show a cement stratigraphy in the sparitic areas, indicating void filling by a sequence of cement precipitation. The CL views show variation in cement history, with bright and dull luminescent cements occurring at different stages in the history. A common interpretation is that bright cement represents early burial low-oxygen conditions where bacterial sulfate reduction (BSR) removes iron from the porewaters precipitated as pyrite, so that manganese causes bright luminescence; later, below the zone of BSR, iron adds to the cement to quench the CL resulting in dull images (Scoffin, 1987). This sequence can be envisaged in Fig. 7F, although Fig. 8 shows a different sequence. Whatever the explanation of the history of cementation, it is difficult to visualize such structures as having resulted from permineralization of sponge tissues essentially because the key issue is the problem of recognizing that the structure was originally sponge tissue.

### Network fabrics

Networks are composed of narrow areas of sparite surrounded by micrite, and appear as two broad types: Rectilinear networks comprising mostly criss-crossing straight lines of sparite with nodes (Figs 3E, F; Fig. 4A-D), that are reasonably interpreted as spiculate sponges; and Curved networks of uncertain origin comprising convoluted curved areas of sparite (Figs 9-12), which vary in structure from those with equal thickness sinusoidal sparite-filled areas to those that are more haphazardly arranged. Both Rectilinear and Curved network types in thin section give the impression that they must exist as a three-dimensional (3D) network (e.g. Luo & Reitner 2014, 3D reconstruction). Rectilinear and Curved networks in some cases resemble opaline spicule networks known from well-preserved Palaeozoic Heteroscleromorphs (lithistids; Figs 3F, 4 and possibly Fig. 9).

**Figure 11.**
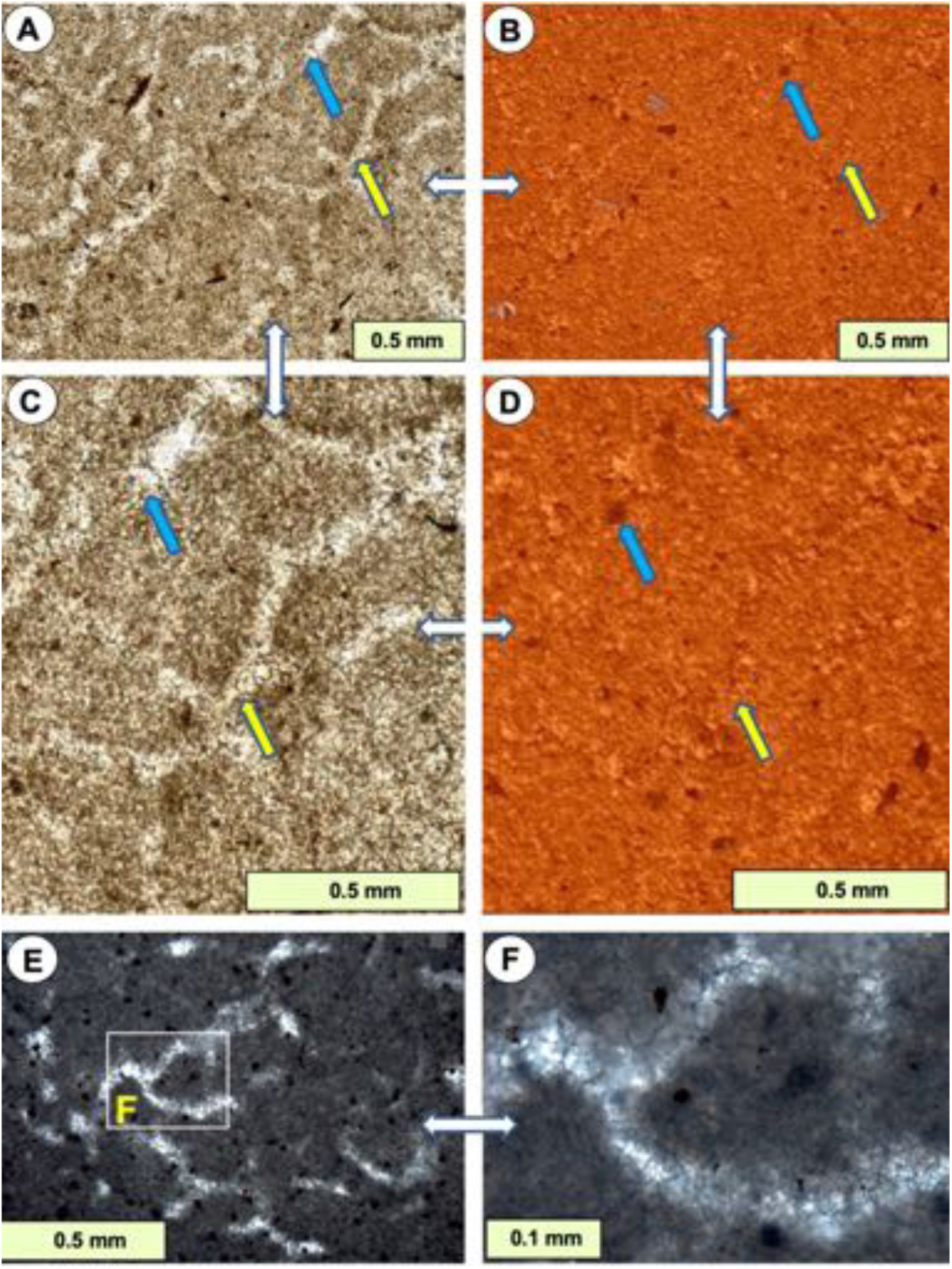
(A-D) Vertical sections of vermicular structure in PPL and CL views. (A) light branched and curved areas are sparite embedded in micrite. (B) CL view of the same area as A. (C, D) Enlargements of A and B respectively; arrows show matched points in the four photographs. The CL view (B, D) shows little difference in luminescence pattern between the two components; the sparite contains poorly luminescent and bright luminescent areas, and the micrite shows a similar variation at a smaller scale, in fine grained material, giving it a speckled appearance. Some portions of the sparite are indistinguishable in the CL view. Whether this arrangement of PPL and CL patterns supports or denies a keratose sponge origin of the vermicular structure is open to discussion. From the boundary between Diplora Beds and overlying Tarnowice Beds, Muschelkalk, Middle Triassic, Libiąż Quarry, Upper Silesia, Poland. (E, F) Vermicular structure from Neoproterozoic carbonates described by Turner (2021, Extended data Figure 1B-C. Note the sparite-filled network fabric. Reproduced under Creative Commons licence (http://creativecommons.org/licenses/by/4.0/), with acknowledgment to Nature.

**Figure 12.**
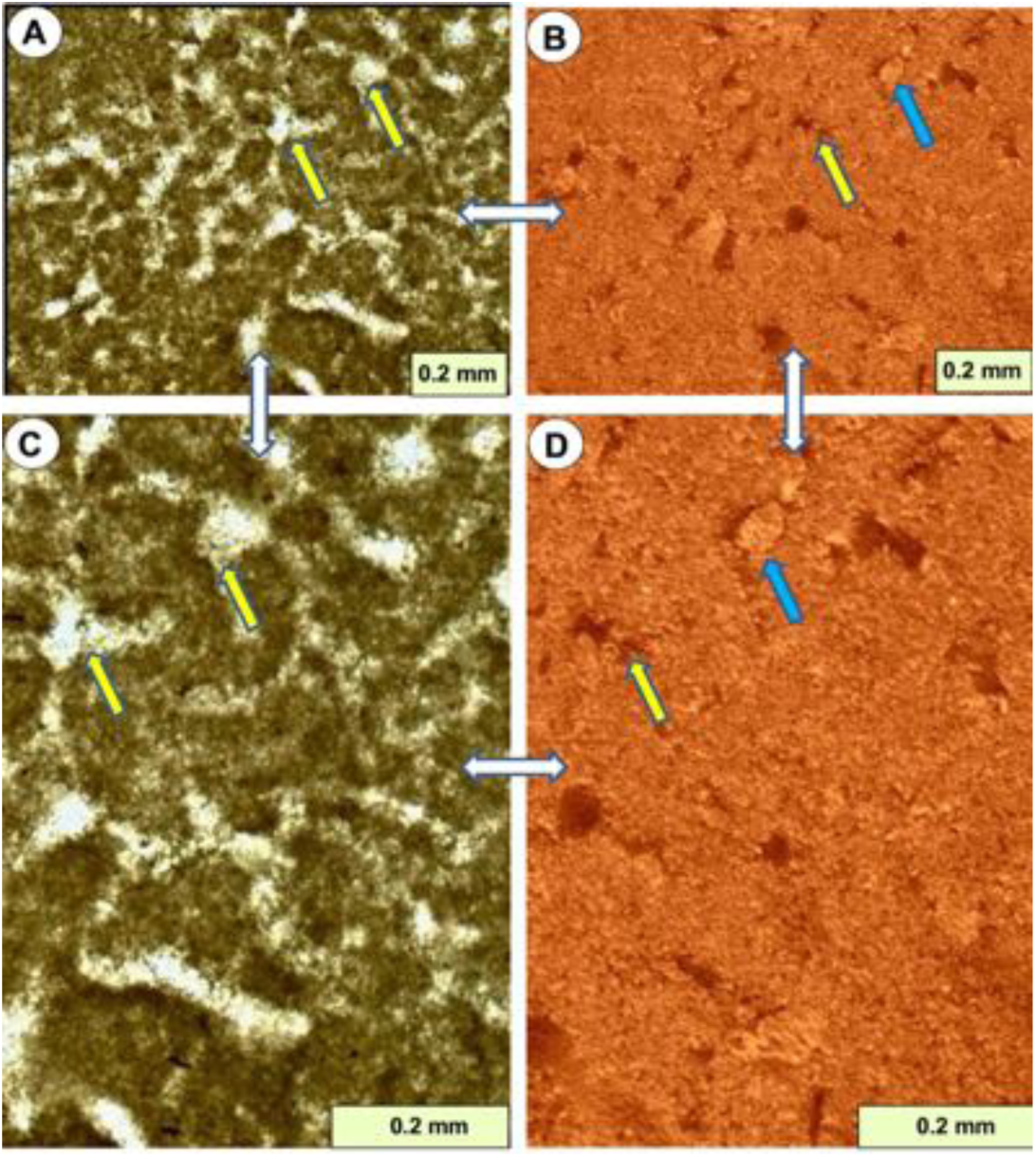
Vertical sections of vermicular structure in PPL and CL views, for comparison with Figs 7, 8 & 11. (A) Light branched and curved areas are sparite embedded in micrite; (B) is the matched CL view. (C, D) Enlargements of A and B respectively; arrows show matched points in the four photographs. Although initial examination indicates differences from Figs 7, 8 & 11, CL patterns in these figures only show variable amounts of poor and bright luminescent areas in the sparite, yet some parts of the sparite are also indistinguishable from the micrite in CL view. Collected from the boundary between Diplora Beds and overlying Tarnowice Beds, Muschelkalk, Middle Triassic, Libiąż Quarry, Upper Silesia, Poland.

Curved networks are illustrated in numerous studies from Neoproterozoic (Fig. 11E, F reproduced from Turner, 2021), Cambrian, Ordovician (e.g. Lee & Hong, 2019) and Permian-Triassic boundary microbialites (Brayard *et al.,* 2011; Friesenbichler *et al*., 2018; Baud *et al*., 2021; Wu *et al*., 2021). Network fabrics found within micrite inside articulated shells, embedded in micritic matrix lacking the nextworks, were interpreted by Park *et al.,* (2017, fig. 3) as spicule networks. Fig. 10 shows examples of curved networks and microbial structures within microbialites directly after the end-Permian extinction; and Figs. 11A-D and 12 explore more variations in Triassic curved networks using both plane light and cathodoluminescence (CL), showing the variation in diagenetic history between the sparite areas and micrite areas. In particular, curved networks may grade into peloidal and amalgamated fabrics described below.

### Amalgamated fabrics

Amalgamated structures comprise patches of micrite, which in some cases give the impression of a vague individuality merged together with intervening spaces occupied by sparite (Fig. 8, which also shows layers, visible in Fig. 8A, B); samples viewed with CL show different cements in the sparite areas compared to the micritic areas, thus indicating voids filled with cement. They overlap with the concept of clotted micrites, but clotting implies a process of sedimentary material sticking together, which may or may not be appropriate in this case, so amalgamated is used here. The possibility exists that some measure of diagenetic change may have affected such material. Amalgamated fabrics have been described as keratose sponges by Lee *et al*. (2014) and Park *et al*. (2017, fig. 3), and were the subject of discussion by Kershaw *et al*. (2021a) in comparison with possible sponges.

### Granular fabrics

The Granular category comprises micritic objects with irregular areas of sparite cement in spaces between objects (Fig. 13). In their simplest appearance they may be described as peloids and commonly occur in cavities forming geopetals (Fig. 14, but compare with Figs. 16 and 17 considered in the discussion). In many of the cases similar to Fig. 14 attributed by authors to keratose sponges (e.g. Lee & Hong 2019, fig. 2; Park *et al*., 2017, fig. 3E, F), the granular fabric grades downwards into the amalgamated fabric, that is, these fabrics give the impression of an evolution of fabric from particulate to amalgamated in function of stratigraphic polarity and packing density. Because disaggregation of peloids into more diffuse masses of micrite is a common phenomenon, careful observation of the intergranular and shelter porosity (thickness variation, grain-supported texture, sagging and dragging along pore walls) holds the key for discrimination of an essentially physical (abiotic) origin.

**Figure 13.**
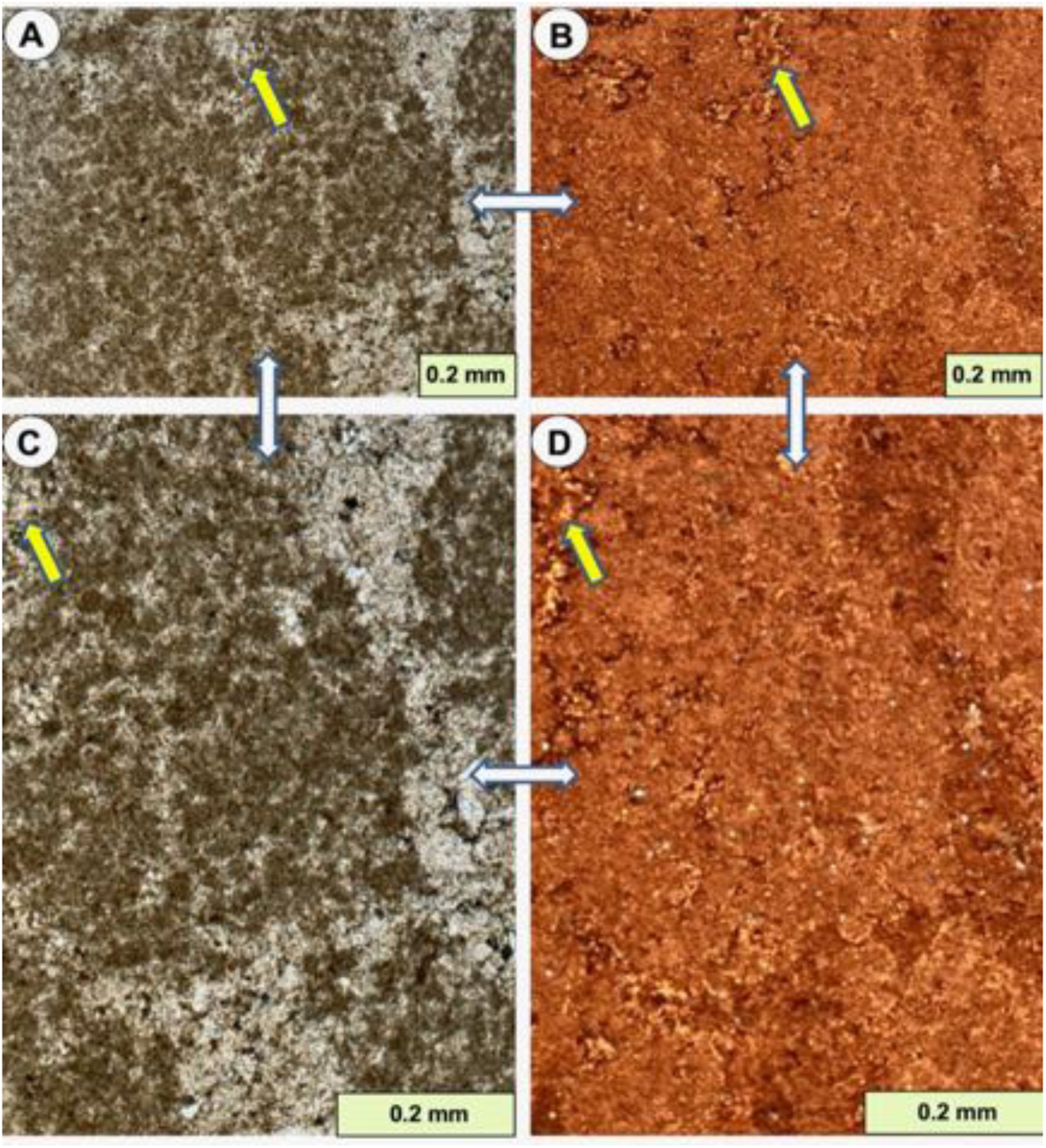
Vertical sections of pelletoidal structures from Viséan limestones of the Boulonnais inlier, N. France. (A, B) PPL (A) and CL (B) views of peloidal carbonate, showing bright orange luminescence of peloids; and dull orange to yellow luminescence of interpeloidal calcite cement. (C, D) Details of the structure, arrows mark matched points. These images are provided here to demonstrate that peloidal fabrics are not compatible with the interpretation as sponges, and may instead be particulate carbonate or microbially deposited; there is no reason to consider these as sponges.

**Figure 14.**
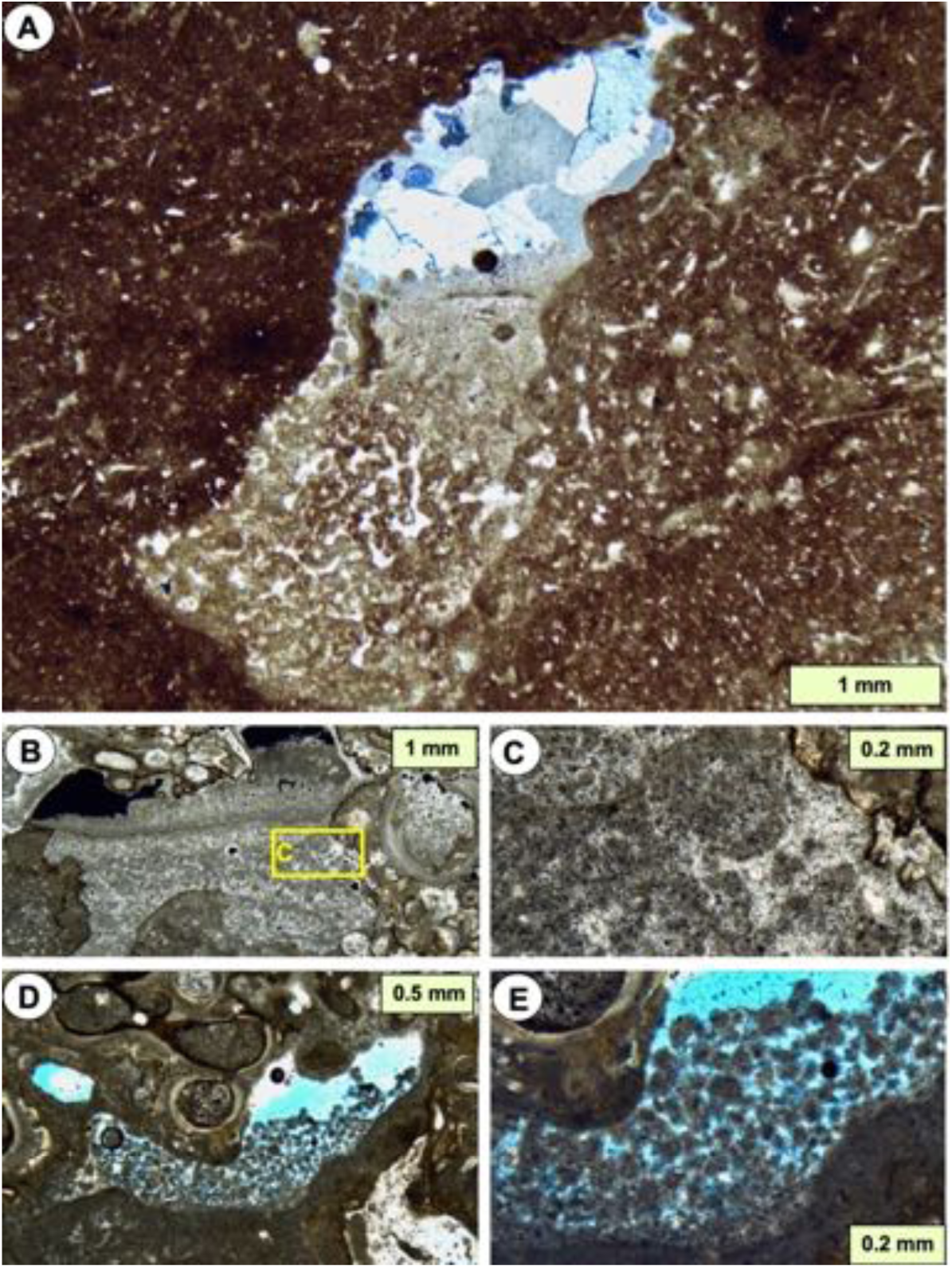
Vertical sections through geopetal fabrics in early-formed cavities in shallow marine limestones, containing peloidal and clotted micrites. (A) Geopetal cavity in a mud-rich coral reef, shows variation from separate peloids at the top down to amalgamated fabrics in the lower part; these were interpreted by Kershaw *et al*. (2021a) to be either inorganic or microbially related structures, in contrast to the interpretation of similar structures as keratose sponges. Huashitou reef, Ningqiang Formation, lower Silurian, Guangyuan, N. Sichuan, China. Reproduced from Kershaw *et al*. (2021a), under CC-BY-NC 4.0, with acknowledgement to Yue Li, The Sedimentary Record and SEPM. (B, C) Geopetal cavity in an algal reef, with layered peloids interpreted as sedimentary, with possible microbial influence. Note that B is in XPL and the two black areas at the top are holes in the thin section; C is in PPL. Late Quaternary, Aci Trezza, eastern Sicily, Italy; after Kershaw (2000). (D, E) Geopetal cavity in algal-coral reef with interpreted particulate peloids and cements. Both images are PPL; blue colour is resin-filled empty space in the geopetal. Holocene, Mavra Litharia, central south coast of Gulf of Corinth, Greece; after Kershaw *et al*. (2005).

**Figure 15.**
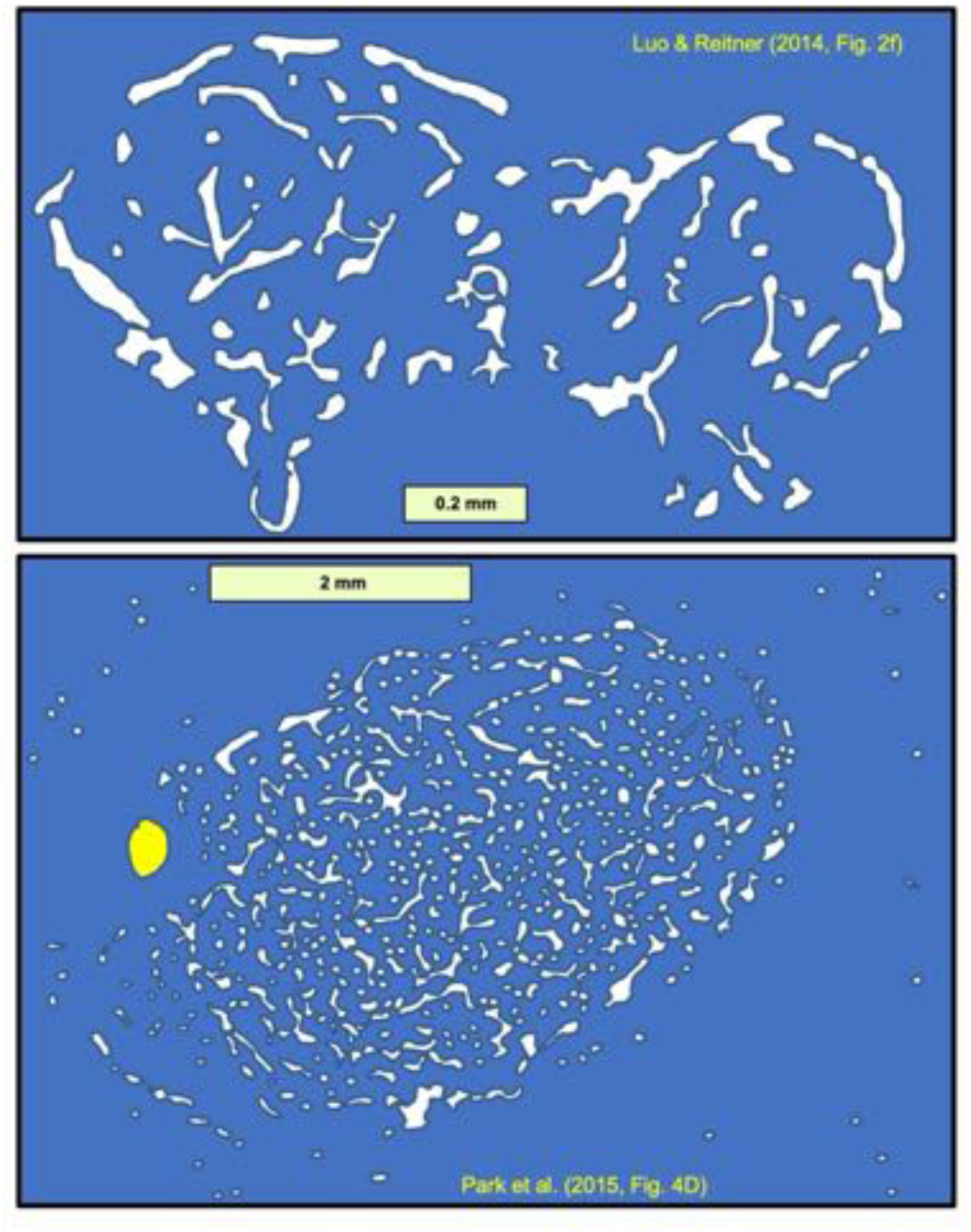
Traced drawings of variegated fabrics, from (A) Luo and Reitner (2014) and (B) Park et al. (2015) to show the pattern of sparite (white), with an outer broken border of curved areas of sparite. The blue background in each case is micrite lacking any clotted or automicrite fabrics and is presumed to be deposited sediment. In B the yellow ellipse is likely an ostracod shell.

**Figure 16.**
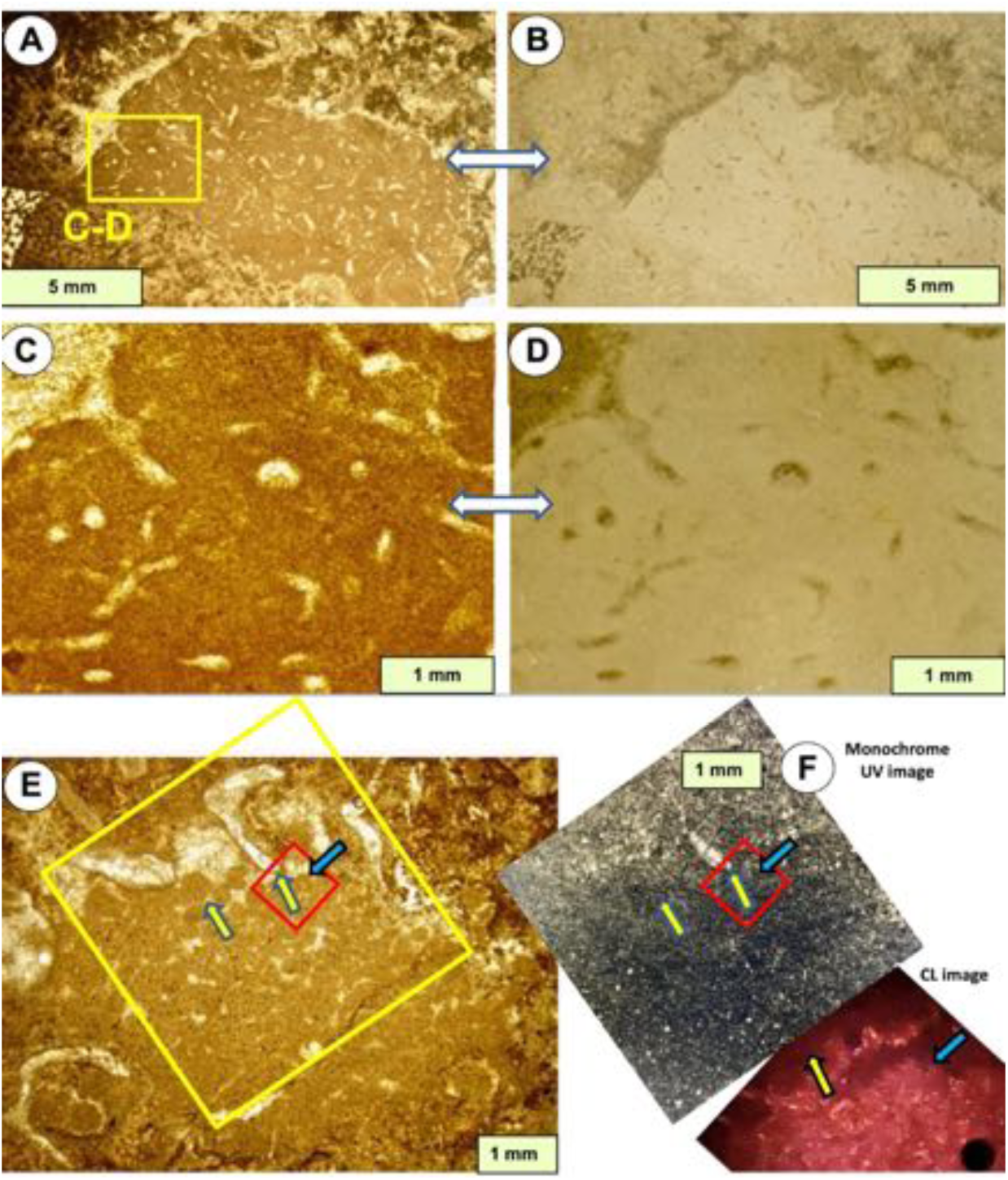
Vertical sections through vermicular structures in cavities in Cambrian carbonates, interpreted by McMenamin (2016) as due to the actions of meiofauna rather than evidence of sponges. (A, B) PPL (A) and reflected light (B) views of vermicular structures within a cavity in an archaocyath-algal boundstone. In this example, the vermicular structures are interpreted as possible graphoglyptid trace fossils comprising microburrow swarms, described by McMenamin (2016). (C, D) Details of box in A, using PPL (C) and reflected light (D) views. Puerto Blanco Formation, Lower Cambrian, base of unit 3, Cerro Rajón, Sonora, México. (E, F) Vermicular structure in the interior space of a dead archaeocyath, here interpreted as comprising packed faecal pellets in a cavity. E is PPL, F shows UV (upper image) and CL (lower image); red box in the UV image shows location of the CL photo; yellow and blue arrows show matched points between these three images. Pellets are discrete in the upper part of the cavity but become more diffuse downwards, interpreted by McMenamin (2016) to indicate pellets disaggregated in the lower part of the pile and the light areas are interpreted as microburrows in the sediment, the burrowing activity may have caused disaggregation of the pellets. Poleta Formation, Cambrian Stage 3, Barrel Springs, Nevada, after McMenamin (2016).

**Figure 17.**
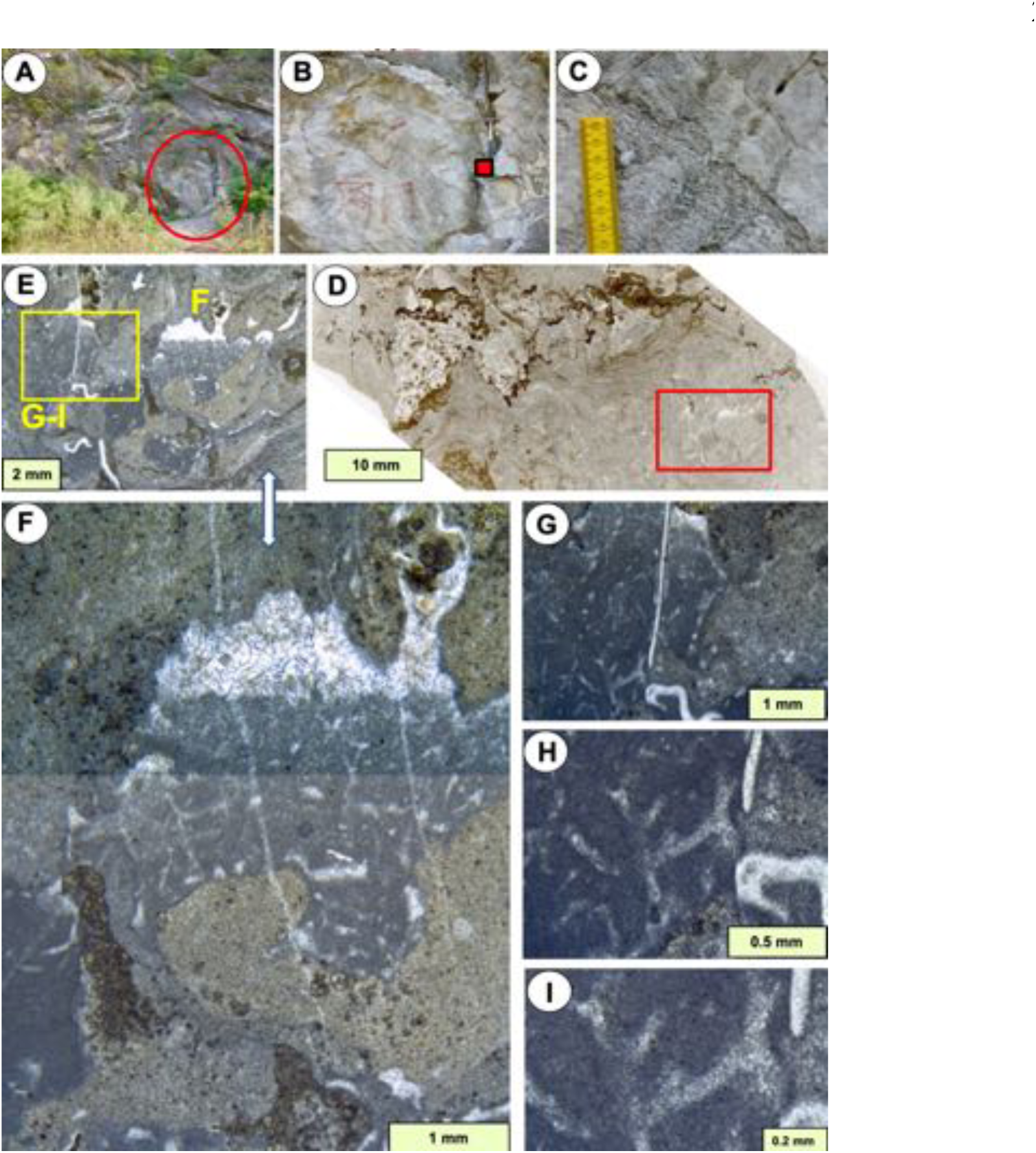
Vertical sections through vermicular structures in a cavity within a stromatolite. (A-C) Field views of a stromatolite bioherm; red box in B shows location of sample, from the upper part of the bioherm; C shows field detail of stromatolite columns overlain by bedded limestone. (D) Whole thin section view of stromatolite, showing abundant cavities in its structure, shown clearly in E (red box). (E) Cavities (darker grey with geopetals) in stromatolite mass. (F) Detail of right side of E, showing vermicular structure in the geopetal fill. (G-I) Details of left side of E (yellow box) showing branched structure of light areas of sparite within the micrite fill of the cavity. These are interpreted here as possible microburrow networks of meiofauna, and not of sponges. Uppermost Gushan Formation, upper Cambrian, Xiaweidian, near Beijing, China.

### Variegated spar fabrics

In some cases attributed to keratose sponges, a separate category of sparite within micrite masses comprises a structure that appears to be organized differently from the Networks described above (Fig. 15, diagrams traced from publications). Variegated structures comprise an outer portion of short lines of sparite that curve round to form the outer limits of a discrete structure, and the inner portion is similar to the Network forms described above. Examples are in Luo & Reitner (2014, fig. 2f), Park *et al*. (2015, fig. 4D; 2017, fig. 4C) and Friesenbichler *et al*. (2018, fig. 10B). These variegated structures are different from the curved networks and are presumed to have been formed by a different process; they seem to occur mostly in cryptic positions, although the case illustrated by Friesenbichler *et al*. (2018, fig. 10B) is in open space between microbialite branches.

## DISCUSSION

The issue expressed in this study is that structures considered to be keratose sponges by numerous authors are unverified, and are even unlikely because of the preservation issues. Thus, such features can be interpreted as other structures, as indicated in the Results section and discussed below. There are four principal areas of concern: 1) verification of the keratose (aspiculate) sponge affinity; 2) alternatives to sponges; 3) accuracy of reporting; and 4) the impact on understanding of ancient ecosystems. One of the prominent difficulties in assessing published illustrations is the low resolution of images, and the common use of thick microscope sections that lack clarity.

### Verification of keratose sponge affinity

The wide variety of fabrics attributed to keratose sponges in the ancient record suffers from lack of verification and coherence, and in the case of the report by Turner (2021) of an early Neoproterozoic example (Fig. 11E,F), the age predates significantly the time corridor predicted for keratose sponges by molecular phylogeny (Fig. 5). In few cases are a mesoscopic body or overall shape reported. Even at the microscopic scale, there is a fundamental problem because no mineralized sponge fabrics are certainly identified, so that preservation of the purported spongin skeleton requires understanding of a diagenetic pathway that seems to have no equivalent in the rock record.

Overall, the claim for fossil keratose sponges in ancient carbonates requires both a proper identification of sponge structure in concert with an exceptional preservation mechanism. Indeed, relatively decay-resistant structural tissue components such as parts of the extracellular collagenous matrix (ECM), mesoscopic strands and networks of spongin, the various forms of chitin (α, β, γ) and cellulose might get physically preserved or replicated *via* permineralisation or *via* polymerisation (Gupta & Briggs, 2011). For the claim of keratose sponges in carbonates, permineralisation (mummy-style preservation) of the sponge to produce automicrite is considered a prerequisite in order to eventually preserve in 3D a former network of spongin as a calcite-cemented mold. Otherwise, if only the spongin was polymineralised (or permineralised) the result should be severe physical compaction only episodically preserving exceptional details (Burgess-style preservation of sponges; Conway Morris & Whittington, 1985; Butterfield & Nicholas, 1996).

### Sponge mummies, indirect replication of a former spongin skeleton

For a keratose sponge to permineralise it would be necessary to replace the sponge tissue with micrite. Froget (1976, on lithistids) followed by Brachert *et al*. (1987, on hexactinellids) provided examples of Pleistocene to Holocene permineralised (calcified) siliceous sponges. In addition, these authors observed in some detail the concurrent onset of diagenetic alteration of the opaline spicules (dissolution, recrystallisation to calcium carbonate, initial cementation). Reitner (1993, p. 26 and Pl. 4/4) illustrated how a living non-rigid demosponge might preserve its original spicular architecture within automicrite. Neuweiler *et al*. (2007) interpreted an intimate connection of mummification with calcifying organic colloids adsorbed onto and into relatively decay-resistant parts of the former ECM, thus dismantling fibrillar collagen during partial death. However, because spongin is a non-fibrillar collagen (Exposito *et al.,* 1991), during decay, no dismantling into submicroscopic collagen fibers with their associated secondary sorptive attributes (surface area, scaffolding; Neuweiler et al., 2007) is expected to occur. Indeed, permineralising (petrifying) modern sponges, except for being a curiosity, typically are very rich in fibrillar collagen giving them a firm-leathery (e.g. the spiculate *Spheciospongia*, Wiedenmayer 1978) to even cartilaginous consistency (e.g. the petrifying verongimorph *Chondrosia*, Göthel, 1992). Nevertheless, it remains questionable whether that small group of extant sponges is representative of fossil sponge mummies (Neuweiler *et al*., 2007 for full discussion). In many sponges, there is a problem of sheer volume, that is the amount of ECM present in a modern sponge does not match the larger amount of automicrite present in sponge mummies (see *Malumispongium* in Bourque & Gignac, 1983; Neuweiler *et al*., 2007), therefore unresolved microbial-organochemical reactions might be involved, and even dissolved pore-water silica might play a role if opaline spicules were originally abundant (Lakshtanov & Stipp, 2010). Thus if the sparite portions of a vermicular structure represent the spongin of a keratose sponge, then transformation from spongin to sparitic calcite (with perhaps an intermediate step not preserved) would have to occur ***after*** conversion of the intervening soft tissue to micrite to prevent compression of the spongin network in burial.

It should be noted here that the spar-micrite structures illustrated in Luo & Reitner (2014, 2016), Lee & Riding (2021a, b) and Turner (2021), in concert with our own results, do not show major compression. Automicrite (mummification) is stated as being present, but no supporting petrographic or geochemical evidence is provided (parameters include: gravity-defying, secondary porosity, fragmentation, local collapse, fluorescence, intracrystalline organic compounds; Neuweiler *et al*., 2000). Another example of the problem of verification is shown in Heindel *et al*. (2018, Figs. 9D, 10B, D), who illustrated microbialites from the well-known Çürük Dag site in southern Turkey. Heindel *et al*. labelled sponges as being present in the matrix that sits between microbial branches, but close examination of those images reveals a calcareous mudstone with minute bioclasts and cannot be considered a sponge mummy. Other images in the same paper show areas of matrix containing fine sparite between microbial branches that may be networks, but there is no demonstration of criteria to indicate that these are sponges mummies; a similar example from south China was discussed by Kershaw *et al*. (2021a).

### Discrete replication of a spongin skeleton

It is conceivable that the spongin itself might be replicated via polymineralization or permineralization, but then preservation of an organic phase (plus compression) or ghost-structures within a permineralizing phase would be expected (Gupta & Briggs, 2011). As noted above, in opposition to the fibrillar collagen in sponge ECM, spongin is a non-fibrillar collagen (Exposito *et al.,* 1991) and during decay or partial death, no dismantling or enhanced secondary sorptive attributes supporting permineralisation (Neuweiler et al., 2007) are envisaged. Another option is coating, that is mineral precipitation and growth at and from the spongin surface (see Szatkowski et al., 2018), but no respective fabric relationship has been reported. The CL images presented in Figs. 7, 8, 11-13 largely show the sparite portions to have different cements from the micrite; in some cases (e.g. Fig. 7F, 8, 12) there are zoned cements in the sparite that indicate early porosity and permeability, so this seems to preclude any mineralization process related to the spongin itself, at least for these samples. Even in Fig. 11A-D, where the distinction between the micrite and sparite areas in CL is minimal, parts of the sparite show different CL response from the micrite. Thus, in the cases illustrated in this paper, there is no obvious basis for mineralization of the spongin itself to explain the sparitic areas. It is easier to explain the CL images in terms of various kinds of early and fabric-selective porosity. Nevertheless, this does not necessarily deny a sponge affinity but leaves mummification as the only option left to explain preservation of keratose sponges in carbonate rocks. However, as stated earlier, the petrographic or geochemical evidence for mummification is either unclear or absent. Finally, there are sponges which contain opaline spicules attached to prominent strands of spongin (Axinellidae) which as fossils should show both spicules together with their associated replication of spongin. No published report of such an intimate relationship preserved in carbonate rock thin-sections was found in the literature.

### Other issues

A significant aspect of observation in relation to sponge affinity in thin-section relates to the pore- and canal system specific to sponges (Figs 1A, B; 2C, D). If sponge mummies are present, a canal system might be preserved in astonishing detail (Neuweiler et al., 2007; 2009) even in the absence of a spicular skeleton (Bourque and Gignac, 1983; Neuweiler et al., 2007, their Fig. 1 A, B). Indeed, Aragonés & Leys (2022) proposed a model for fossil sponge recognition based on the presence of a canal system. However, there are no respective features in all the fossil examples illustrated here and in publications examined in this study. Neuweiler et al. (2009) denied the presence of sponges in early Neoproterozoic polymuds because of the lack of any signs of a preserved canal system.

On the other hand, the canal system (together with spicules) might be too tiny to be visually replicated, although other observations (automicrite, context, substrate) may indicate a sponge interpretation (Shen & Neuweiler, 2018). Lee & Riding’s (2020) reconsideration of the enigmatic structure *Cryptozoön* provides an excellent example of the overall problem of sponge recognition. The fabric interpreted as keratose sponge in Lee & Riding’s (2020, fig. 5c, d) looks somewhat different from the supposed keratose standard image in their fig. 9c, but instead resembles, but not fully matches, the lithistid in their fig. 9a. Despite high quality preservation, there is no (hierarchical) canal system, there is no cortical architecture and there is no analysis of diagenesis. Evidence for the presence of keratose sponges (in opposition to conventional (microbial) spongiostromata) is needed to test the original interpretation (Luo & Reitner 2014, 2016).

In summary (see also Table S1), without verification the presence of fossil keratose sponges in thin-sections made from limestones-dolostones is called into question. Table S1 represents an effort to requalify the most prominent examples as: essentially microbial (spongiostromate, birdseyes-vugular-fenestral porosity), biogenic-problematic, and dubio-to even pseudofossil in nature.

### Alternatives to sponges

#### Endobenthos

Geopetal cavities in lithified limestone and in articulated shells show a common pattern where the upper part of the deposit comprises peloids, that grade downwards into amalgamated micrite within the cavity, as noted earlier (e.g., Fig. 14). Several of these examples are reported as sponges (Lee & Hong, 2019, fig. 2) but they are easily recognizable as peloidal micrites that merged downwards to form clotted structures. The formation process of such features is not obvious. Although they may be considered as reflecting compaction in the sediment mass, it is notable that compaction requires sufficient mass of material to enable gravitational compression, which seems unlikely in such small structures. An alternative is that they may reflect small-organism activity in the cavities, and thus could be meiofauna. The concept of meiofauna (organisms of sizes between micro- and macro-fauna, up to 1 mm size) is well-developed in biological literature (Semprucci & Sandulli, 2020) but almost unknown in the ancient record (e.g. Knaust, 2010). The ability of meiofaunas in modern environments to create burrows and microborings provides a viable alternative to at least some of the possible keratose sponge interpretations described in this study.

Micro-organismic activity is proposed to explain some carbonate facies (McMenamin, 2016), with common occurrence in protected locations such as cavities and empty shells lying on the ancient seabed. Fig. 16 shows a Cambrian example of potential microboring networks in a cavity, noting that the images also indicate geopetal sediment in the sparite areas indicating an open network prior to cementation. Fig. 16E, F explores the use of UV fluorescence microscopy and shows in this monochrome image that the brighter areas (therefore presumably containing more organic matter) are outside the area of the network; this is interpreted to indicate that the network was not composed of automicrite and thus not related to sponge degradation. The accompanying CL image (Fig. 16F) indicates brighter luminescence in the sediment that may reflect diagenetic alteration. Fig. 17 shows a case of cavities inside the outer portion of a stromatolitic dome in shallow marine platform carbonates from North China; the cavities contain micrite and some type of network that does not resemble a sponge, and may be interpreted as a microboring net. In the view of the authors of this study, the examples of peloidal and network structures found in cavities and shells cited above are open to be reinterpreted as meiofauna (microscopic faunas) rather than sponges.

Verification of evidence of ancient meiofauna in sedimentary rocks is in early development (Dirk Knaust, Pers. Comm. 2021; see also McIlroy, 2022). Meiobenthic trace fossils are a relatively new field of ichnology. In the small number of available publications (e.g. Knaust, 2007), foraminifers, nematodes, annelids (particularly polychaetes), arthropods (ostracodes, malacostracans) are listed as the most plausible producers, sometimes being preserved at the end of the trace (Knaust, 2007). Meiofauna burrows may be identified by their constant diameter and regular winding to sinusoidal character, features that are seen in some published photos interpreted by some authors as keratose sponges (e.g., Park *et al*., 2015, fig. 8A), and may explain the fabrics reproduced here in diagram form in Fig. 15). Knaust (2007) felt confident to name some cases as trace fossil ichnotaxa, e.g. *Cochlichnus* Hitchcock 1858. Meiofauna burrows are expected to concentrate in organic-rich areas of the sediment, such as within macrofauna burrows or whole shells; in this context the features in Luo & Reitner (2014, fig. 2f) and Park *et al*. (2015, fig. 4D), reproduced in Fig. 15, are potential macrofaunal burrows penetrated by meiofauna, whereas the figure Park *et al*. (2015, fig. 8B) presents a whole brachiopod shell that may have been passively filled with micrite and subsequently penetrated by meiofauna to produce vermicular-structured micrite.

#### Dubio- to Pseudofossils

3D spar-micrite micro-networks might result from cementation of interparticle porosity of fine-grained granular-pelletoidal sediment material, an important source of ambiguity of carbonate rock petrography (Macintyre, 1985; Lokier & Al Juanabi, 2016; Kershaw *et al.,* 2021a). The issue is complicated because the initial state and cohesiveness of peloidal material varies greatly from loose aggregates-floccules to indurated grains via an entire spectrum of plasticity (Schieber *et al*., 2013). The consequences might be severe because during consolidation and physical compaction the initial granular texture might be lost, resulting in a grumelous ghost structure or even a diagenetic mudstone texture (Lokier & Al Juanabi, 2016 for full discussion). Peloidal textures might also result from authigenesis (automicrite) and heterogenous aggrading neomorphism (Bathurst, 1975; Dickson, 1978; Macintyre, 1985). The examples of peloidal textures in geopetal infills in cavities presented by Lee & Hong (2019) as sponges can be alternatively interpreted as peloidal fills in cavities. In another example, the microspar groundmass in Turner’s (2021) study of Neoproterozoic vermiform structure contains no features that would indicate it originated through ‘permineralization of a pre-existing biological substance’ (automicrite). The illustration of a vermiform microstructure in a shelter void (Turner 2021, extended data, fig. 2) grades into the underlying and overlying homogenous microspar; the lack of a sharp contact between the vermiform area and adjacent micrite reduces confidence that this structure is a sponge. The Early Neoproterozoic vermiform microstructure (Turner, 2021) may indeed be a dubiofossil or even pseudofossil, possibly caused by fluid escape during the consolidation of a flocculated gel-like carbonate mud (syneresis). Fluid escape, volume loss and microfolding (Turner 2021, extended data, fig. 2) raises the possibility of relationship with molar tooth structures of the Neoproterozoic (carbonate gels of Hofmann, 1985 for the Little Dal Group; Kuang, 2014 for review). Furthermore, although they have some resemblance to microburrow nests observed in Phanerozoic limestones (graphoglyptid trace fossils, see Kris & McMenamin, 2021), it seems unlikely that endofaunal metazoans would have existed in the Early Neoproterozoic, so the respective claim for presence of a worm-like (bilateralian) organism (Kris & McMenamin, 2021) would even intensify the conflict with respect to molecular-clock divergence-time estimates.

### Accuracy of reporting

As stated earlier, examination of literature on keratose sponges in ancient carbonates reveals two common features: 1) that all studies refer back to the original 3D reconstruction study by Luo & Reitner (2014); and 2) that in almost all cases, subsequent authors regarded these structures as actual sponges without further investigation. Also, there are cases of misreporting earlier studies, giving the impression of occurrence of sponges in other sequences, that were **not** stated in the cited works; this has resulted in cases of inaccurate reporting of possible sponges, without verifying the original sources. A good example of misreporting may be found in literature on the Permian-Triassic boundary microbialite (PTBM) sequences. Ezaki *et al*. (2008, fig. 8C) illustrated a fabric they described as a “Highly amalgamated and interconnected areas exhibiting spongelike texture with infilling of peloids”. This paper was cited by Friesenbichler *et al*. (2018, p. 654) as an example of sponges in the PTBMs, and subsequently included in the compilation by Lee & Riding (2021a, table 1) as keratose sponges. However, careful examination of the material illustrated by Ezaki *et al*. (2008, fig. 8) shows that all four images in that figure are actually partially altered portions of the lobate microbialite constructor of post-extinction microbialites in South China, now named as *Calcilobes wangshenghaii* (partly illustrated in Fig. 10G; also see Kershaw *et al*., 2021b), thus not a sponge. Another example may be found in Heindel *et al*. (2018) who described “possible keratose sponges”, but these were referred to as “keratose sponges” in Lee & Riding (2021a, table 1). Examination of illustrations by Heindel *et al*. (2018), noted earlier, shows that some of those illustrated are simply carbonate mudstones-wackestones. Thus, the notion of a “sponge takeover” after the end-Permian extinction, envisioned by Baud *et al*. (2021), which uses the work of Ezaki *et al*. (2008), Foster *et al*. (2019) and Friesenbichler *et al*. (2018) as examples of sponges in the microbialite, may be premature.

### Implications: the four settings

In the introduction section, four settings of interpreted keratose sponges were presented, and broader aspects of each are stated below to provide perspective of the implications of this study.

1. Neoproterozoic possible keratose sponges were proposed by Turner (2021), therefore indicating possible metazoans at 890 Ma, significantly earlier than the first appearance of possible metazoans in late Ediacaran Period (ca. 575 Ma, see Wood, 2016). Turner’s proposal is therefore potentially highly significant, but relies on verification.
2. Although the concept of consortia between keratose sponges and microbial structures is proposed (Lee & Riding, 2021a; Pei *et al*., 2021a, b), there are no modern records of keratose sponges as consortia with other organisms in normal marine environments. Nevertheless, Ellison *et al*. (1996) demonstrated an unusual mutualism between sponges and roots of mangroves in shallow subtidal oligotrophic settings in Belize, serving as a reminder of the enormous ability of sponges. Furthermore, it is important to recognize that sponges have copious assemblages of bacteria in their tissues, so it does not preclude the possibility of consortia in ancient times, but the lack of reported consortia between sponges and stromatolites in modern environments thus means that no modern analogues for the ancient carbonates have yet been found.
3. Cambro-Ordovician occurrences of potential keratose sponges reported in numerous studies have importance for Palaeozoic evolution of the biosphere with impact on understanding the Great Ordovician Biodiversity Event, noted by Servais *et al*. (2021) to consist of an episode of change rather than a short-term event. If sponges occurred in larger abundance than has been recorded by verified sponges, then there is an important potential impact on the nature of ancient benthic assemblages across this period. Thus, it is critical to correctly identify the affinity of these structures before applying them in a wider context of biodiversity.
4. Related to point 3, rapid and immense shifts in ecosystems after the end-Permian extinction include a short period of development of microbialites in shallow marine carbonate settings, the appearance and disappearance of which have not been explained. However, recent contributions to literature of interpreted presence of keratose sponges has a significant impact on models of biotic and environmental change, so correct identification is critical. The Permian-Triassic boundary microbialites are likely unique in the rock record (Kershaw *et al*., 2021b), so the notion of a concurrent sponge increase is enormously potentially influential in ecosystem analysis. Furthermore, interpretation of keratose sponge expansion after the end-Permian mass extinction is an attractive idea, corresponding with the notion of sponge development during the low-oxygen conditions associated with that extinction, because sponges are known to tolerate low oxygen conditions. Nevertheless, the lack of sponges in modern stromatolites, and lack of verification of sponges in post-extinction facies means that it is not wise to include sponges in models. The corollary is that the unverified reports of sponge presence also lead to uncertainty in the nature of biotic assemblages. In a modern context, there is increase in sponges in modern coral reef systems that may be a reflection in the decline of corals, while sponges are more resilient to change. Sponge expansion after mass extinction is thus an area of great potential interest in understanding modern changes but needs to be verified. A key point in this debate is that modern living sponges normally disintegrate and disappear from the biota (Debrenne, 1999) in a very short period after death, so from the point of view of ‘the present is the key to the past’, there is a problem of determining abundance and diversity of sponges through their geological history, due to poor overall preservation potential.

In summary, none of the reported examples of keratose sponges in ancient limestones are supported by criteria, and in some cases, sponges are demonstrably absent. This does not necessarily mean that none of the others are keratose sponges, but the lack of proven sponges now requires a concerted effort of objective science to sort out this problem and prevent this snowball of uncertainty continuing to grow.

### CONCLUSIONS

Key points emerging from this study are:

1. The interpretation of keratose sponges (that consist of skeletons lacking mineral components) in carbonate facies through the Neoproterozoic to Triassic record (and likely the entire geological record) may or may not be real, with implications for the palaeobiology and evolution of sponges, and palaeoecology of fossil assemblages. The interpretation is considered here to be at best unsafe, and at worst incorrect, so the importance of keratose sponges in geological history remains uncertain.
2. The problem of how a keratose sponge spongin network may come to be preserved as sparitic calcite is a critical and unexplained component of diagenesis that needs to be addressed.
3. All published studies claiming keratose sponges need to be re-examined to confirm or deny their presence; such work may overturn current ideas of the role of keratose sponges.

## ACKNOWLEDGEMENTS

During development of this study we are very grateful for discussions with Andrej Pisera, who made valuable comments on an earlier version of the manuscript, and provided images used in Fig. 3E,F. Comments from Joe Botting, Marcelo Carrera, Cristina Diaz, Dirk Erpenbeck, Jonsung Hong, Dirk Knaust, Jino Park, Christine Schönberg, Elizabeth Turner, Max Wisshak and Gert Wörheide have provided much insight into this study. SK thanks: Haizhou Wang (China University of Petroleum) for access to Cambrian carbonates in North China illustrated in Fig. 17; Yue Li (Nanjing Institute) for access to Silurian carbonates in South China, including images shown in Figs 9 & 14; and Robert Riding (Tennessee) for access to Cambro-Ordovician carbonates in Utah illustrated in Figs 4 & 9. CS also thanks: Lucie Goodayle, Natural History Museum, London, Science Photographer for *Vauxia* images; Tom White and Javier Ignacio Sánchez Almazán, from Natural History Museum, London, and National Museum of Natural Sciences, Madrid, respectively, for access to Recent sponges. We gratefully acknowledge information on microborings provided by Klaudiusz Salamon, who sadly died in early 2022.

## DATA AVAILABILITY STATEMENT

Not applicable

---

**Table S1.**
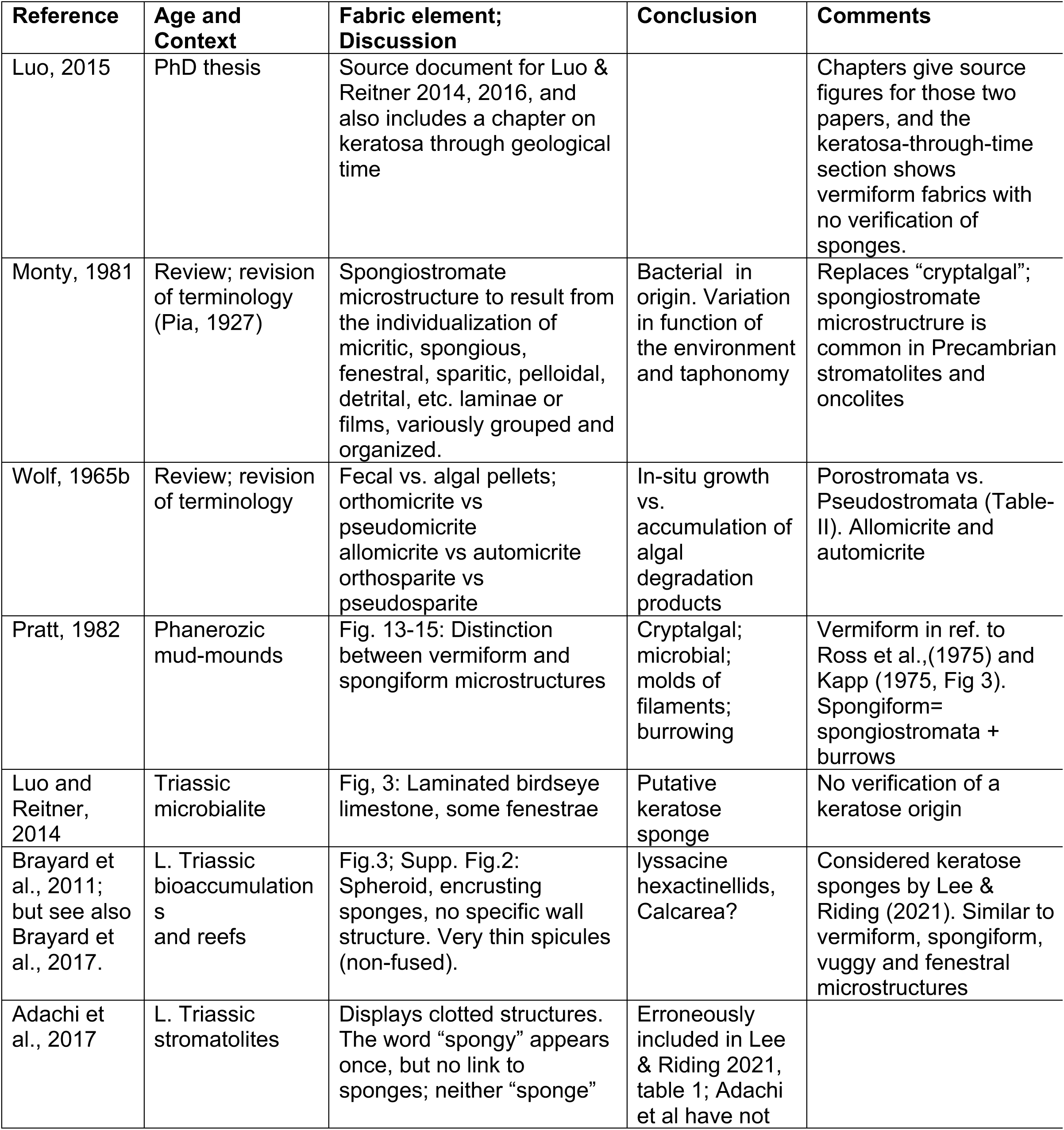
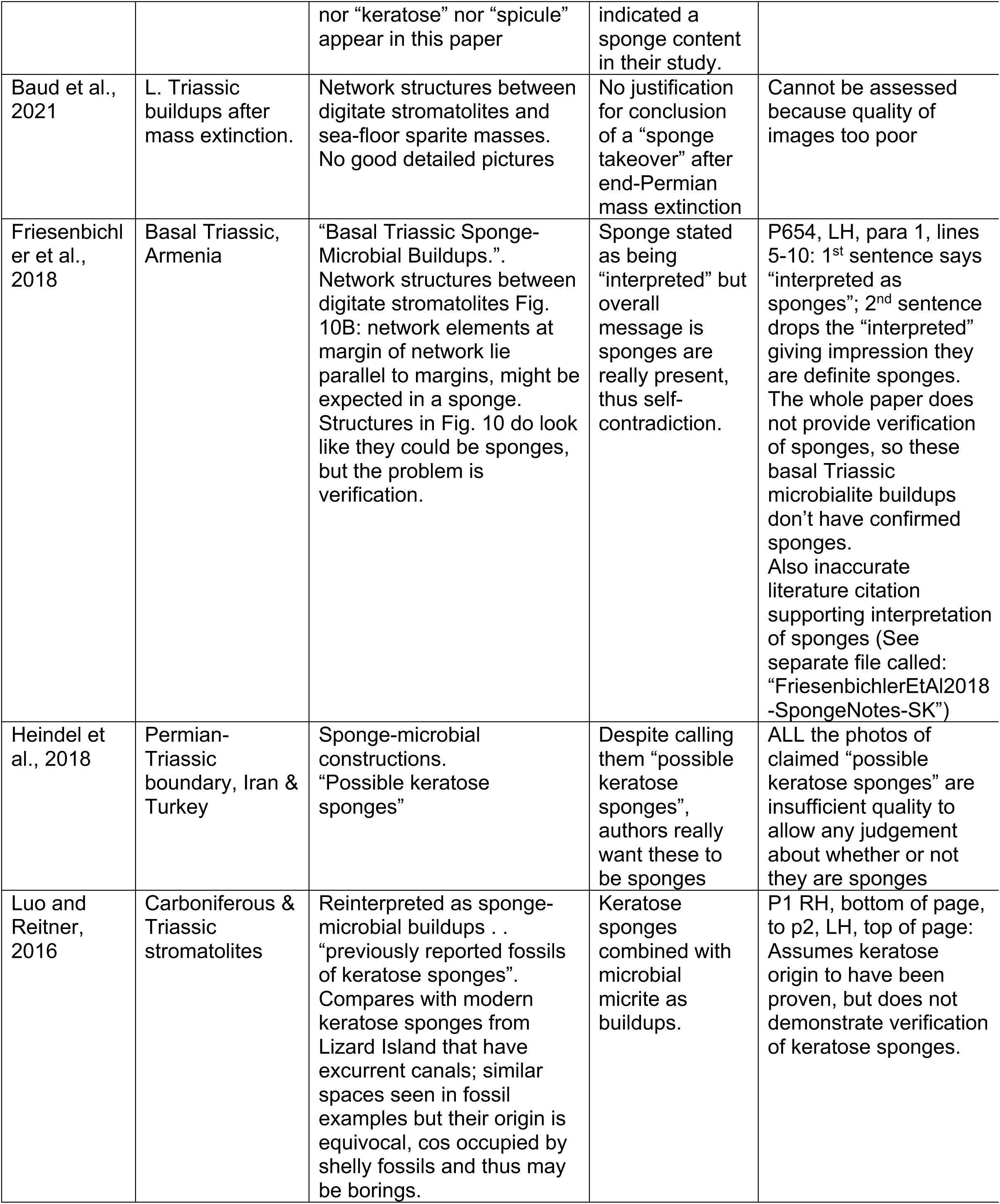
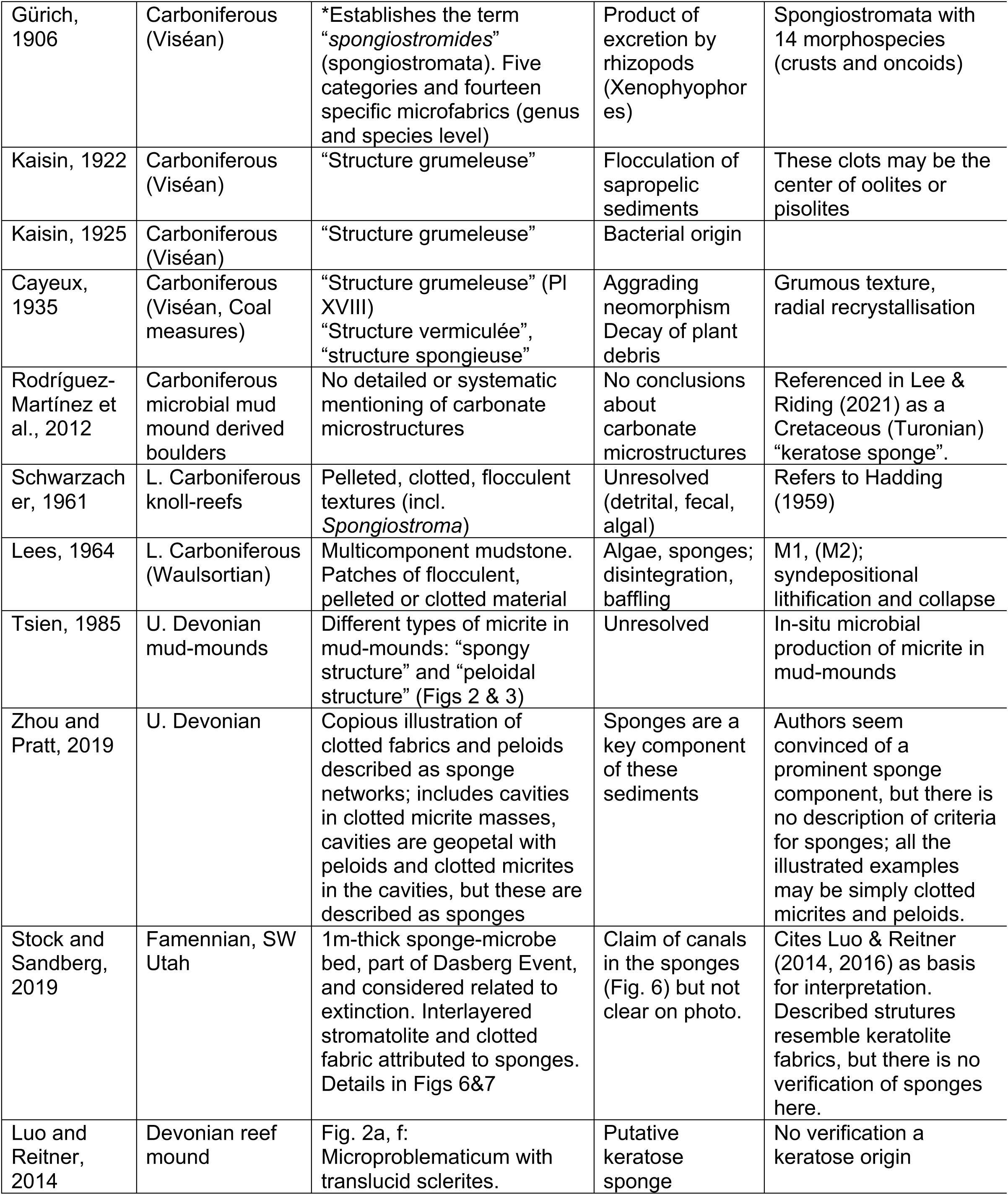
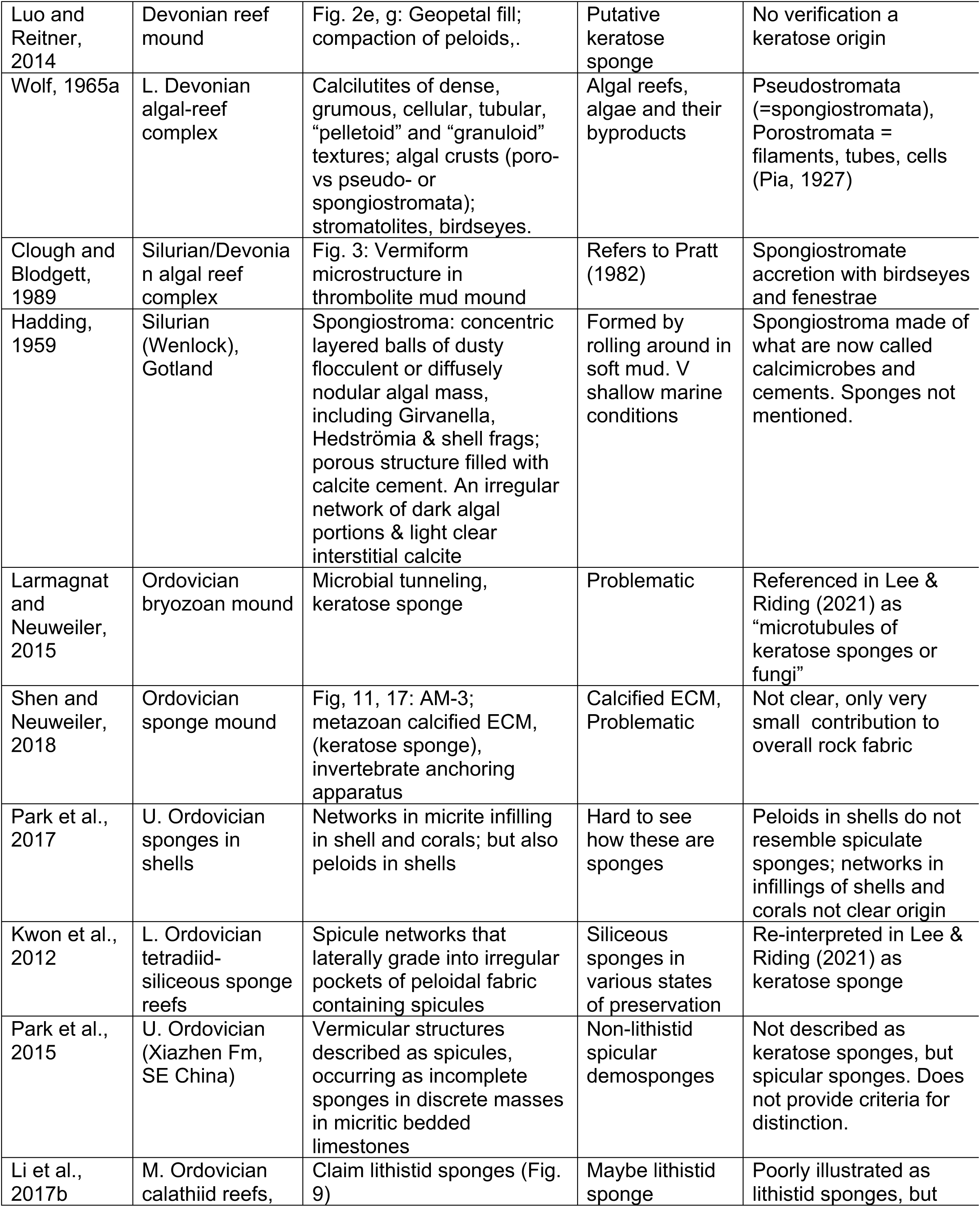
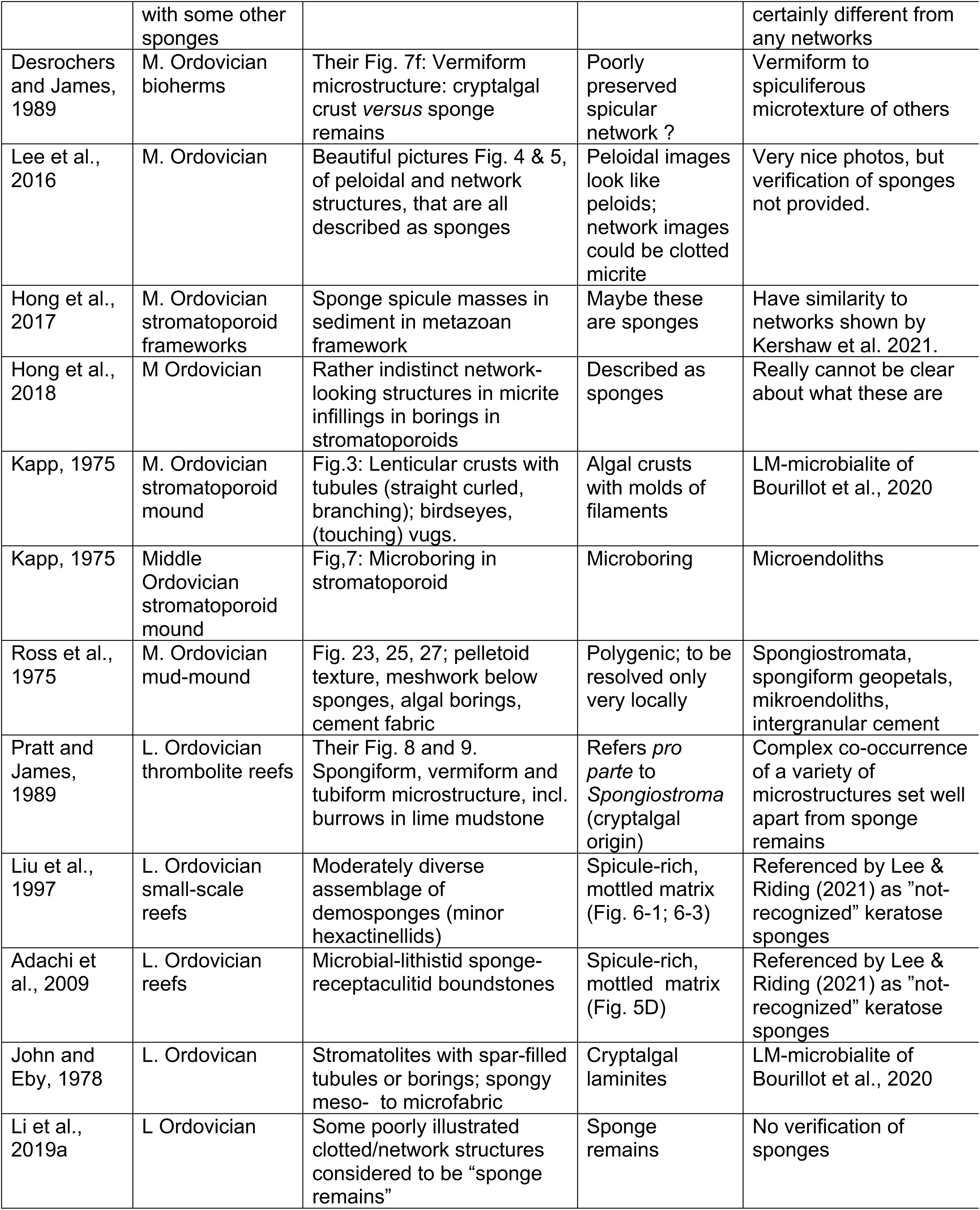
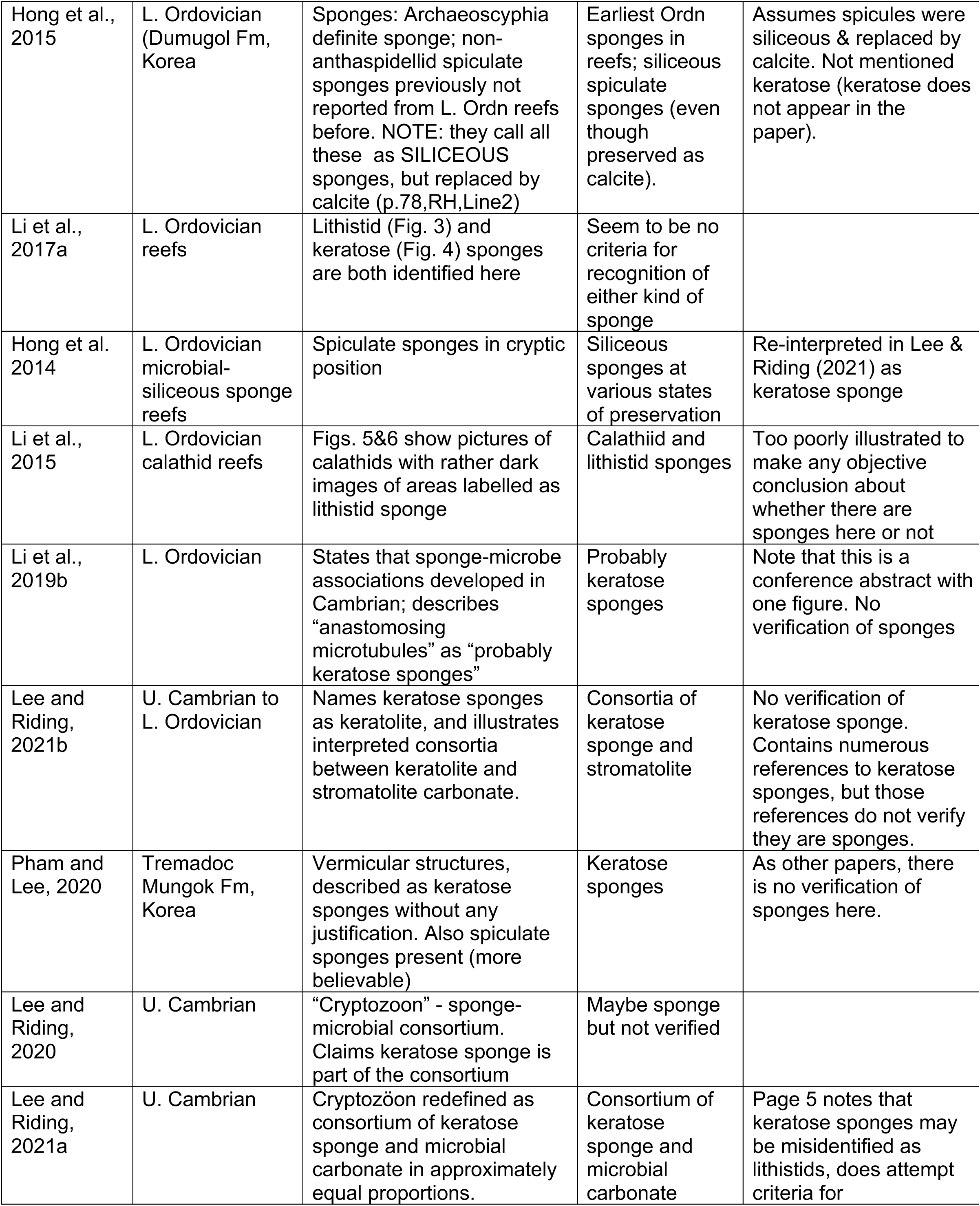
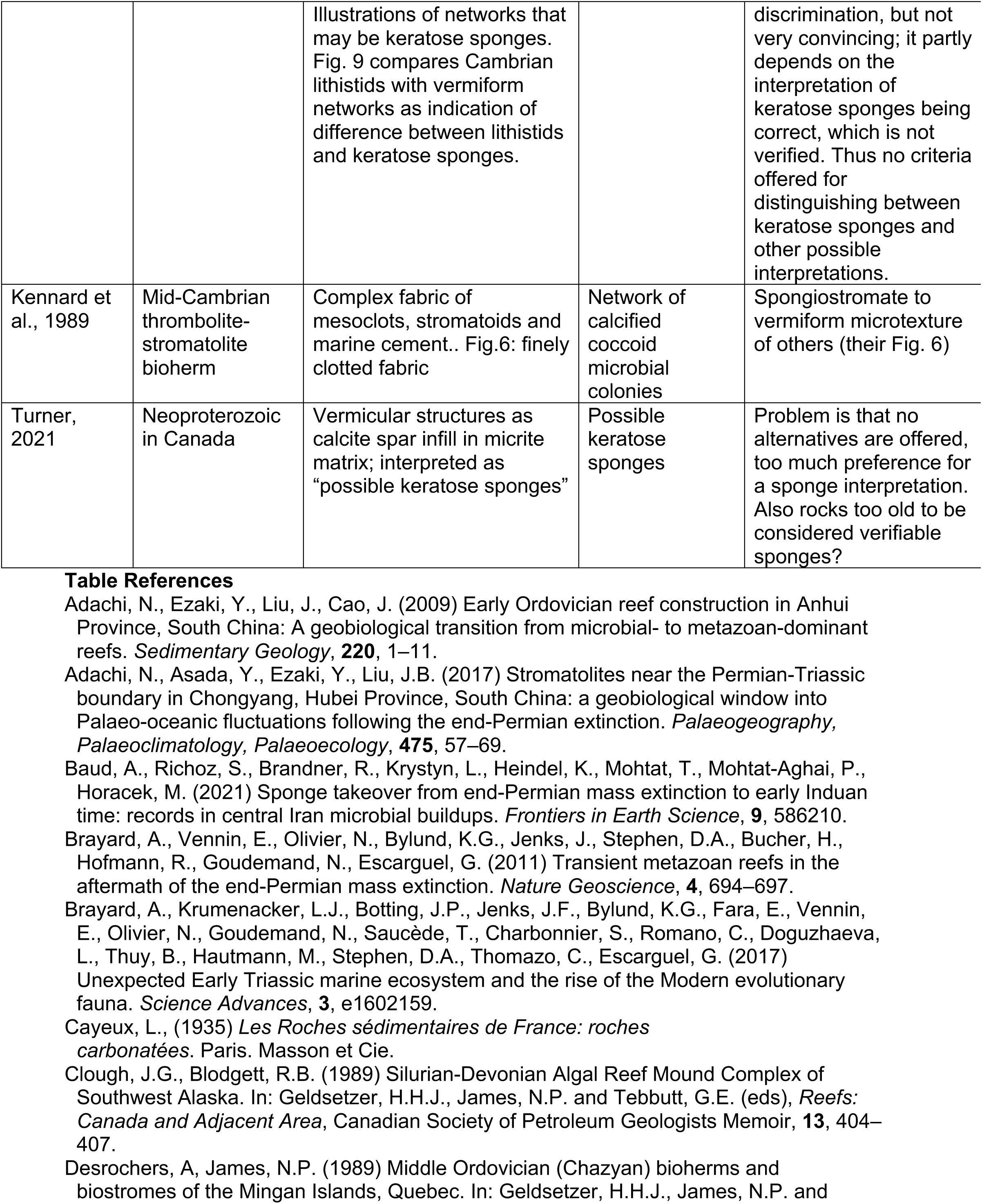
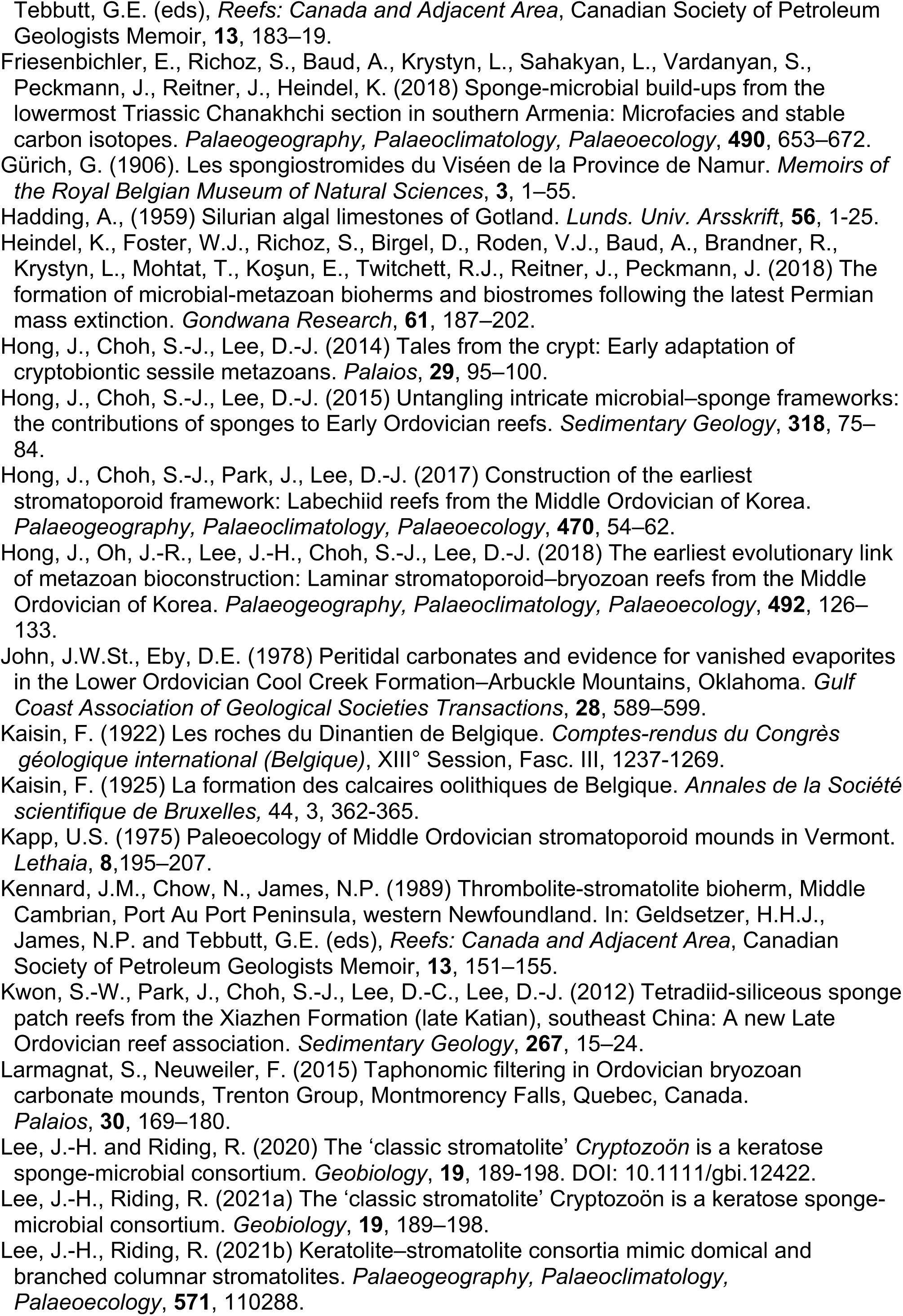
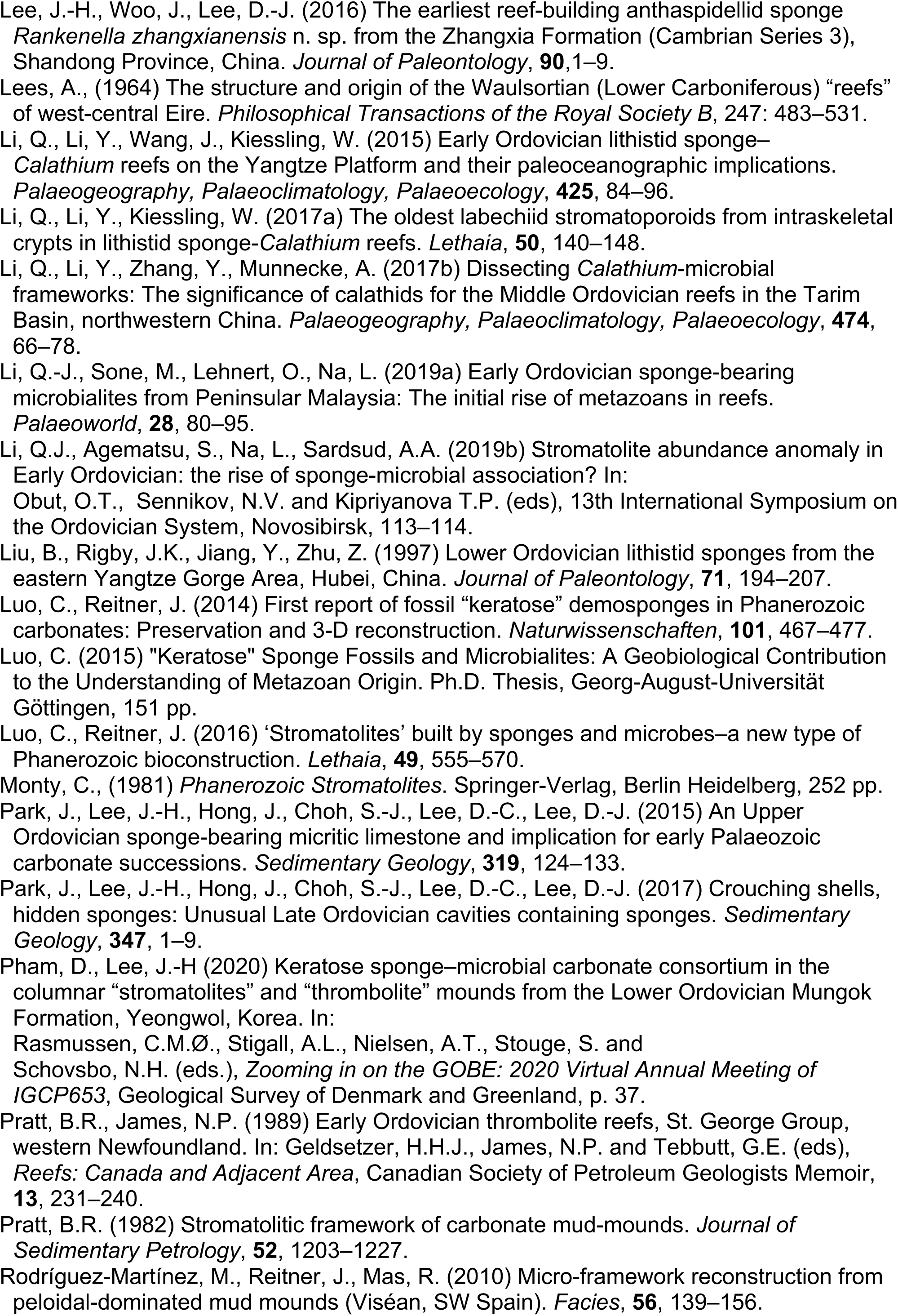
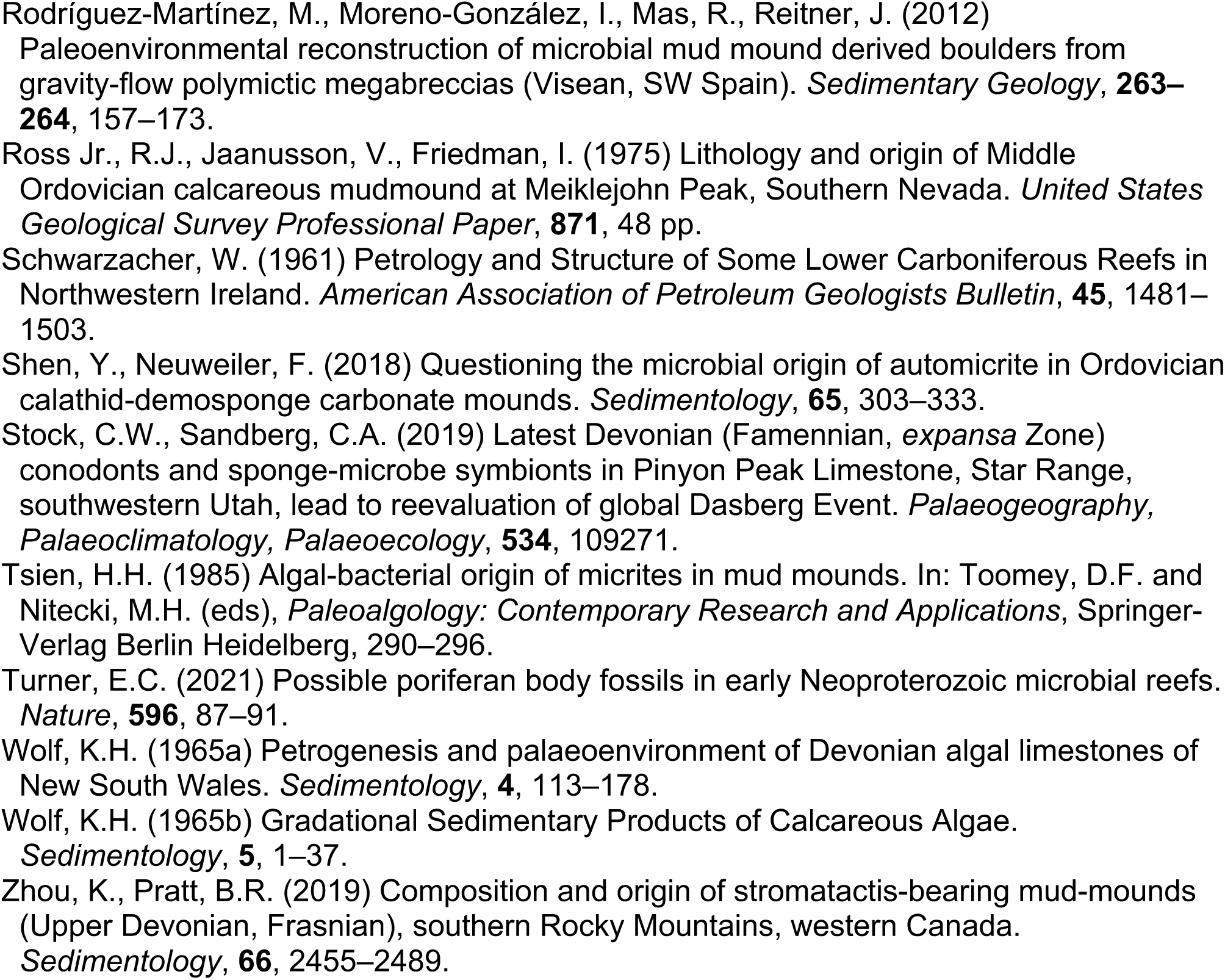
Compilation of publications describing possible keratose sponges, presented in stratigraphic order (oldest at bottom), together with key points and interpretive comments by the authors of this study; the list of references in the table is provided below the table.>>>

